# Resin-acid derivatives bind to multiple sites on the voltage-sensor domain of the Saker channel

**DOI:** 10.1101/2020.06.11.146548

**Authors:** Malin Silverå Ejneby, Arina Gromova, Nina E Ottosson, Stina Borg, Argel Estrada-Mondragón, Samira Yazdi, Panagiotis Apostolakis, Fredrik Elinder, Lucie Delemotte

## Abstract

Voltage-gated potassium (K_V_) channels can be opened by negatively charged resin acids and their derivatives. These resin acids have been proposed to attract the positively charged voltage-sensor helix (S4) toward the extracellular side of the membrane by binding to a pocket located between the lipid-facing extracellular ends of the transmembrane segments S3 and S4. By contrast to this proposed mechanism, neutralization of the top gating charge of the Shaker K_V_ channel increased resin-acid induced opening, suggesting other mechanisms and sites of action. Here we explored the binding of two resin-acid derivatives, Wu50 and Wu161, to the activated/open state of the Shaker K_V_ channel by a combination of in-silico docking, molecular dynamics simulations, and electrophysiology of mutated channels. We identified three potential resin-acid binding sites around S4: (1) The S3/S4 site previously suggested, in which positively charged residues introduced at the top of S4 are critical to keep the compound bound, (2) a site in the cleft between S4 and the pore domain (S4/pore site), in which a tryptophan at the top of S6 and the top gating charge of S4 keeps the compound bound, and (3) a site located on the extracellular side of the voltage-sensor domain, in a cleft formed by S1-S4 (the top-VSD site). The multiple binding sites around S4 and the anticipated helical-screw motion of the helix during activation make the effect of resin-acid derivatives on channel function intricate. The propensity of a specific resin acid to activate and open a voltage-gated channel likely depends on its exact binding dynamics and the types of interactions it can form with the protein in a state-specific manner.

**eTOC Summary:** Silverå Ejneby et al use molecular dynamics simulations and electrophysiology to show that the voltage-gated Shaker potassium channel has multiple binding sites for resin-acid derivatives that can regulate its opening.

## INTRODUCTION

Resin acids, which are primarily found in pine resin, and their chemical derivatives promote the opening of several voltage-gated potassium (K_V_) channels (Ottosson et al., 2015, 2017, 2014; Sakamoto et al., 2017; Salari et al., 2018; Silverå Ejneby et al., 2018) and the voltage-gated and calcium-activated BK channel (Imaizumi et al., 2002; Sakamoto et al., 2006). A naturally occurring resin acid, isopimaric acid, also promotes inactivation of voltage-gated sodium and calcium channels (Salari et al., 2018). One example of resin-acid activated K_V_ channel is the human M-type K_V_ channel (hK_V_7.2/7.3), which regulates the excitability of nerve cells of the brain (Brown and Adams, 1980; Wang et al., 1998). Resin-acid derivatives are therefore interesting drug candidates for the treatment of hyper-excitability related diseases such as epilepsy (Kobayashi et al., 2008; Ottosson et al., 2015; Silverå Ejneby et al., 2018; Wu et al., 2014a).

In contrast to many other channel-targeting compounds, resin acids and their derivatives are relatively hydrophobic (LogP ≈ 6), because of their three-ringed structure. As such, they have been suggested to partition into the lipid bilayer and interact with the *Drosophila* Shaker K_V_ channel from the extracellular membrane-facing side (red triangles in Fig. 1A,B; Ottosson et al., 2017). The negatively charged resin acid is suggested to stabilize the activated state of the voltage-sensor domain (VSD) in which the positively charged S4 helix is in an up state, thereby promoting gate opening in the pore domain of the channel. Another equivalent way to describe this effect is to consider that the resin-acid derivative exerts an upward and clockwise electrostatic pull on S4 (Fig 1C; Ottosson et al., 2014, 2015, 2017; Silverå Ejneby et al., 2018). Thus, by stabilizing activated/open states relative to resting closed states, the resin acids shift the conductance-versus-voltage, *G(V)*, curve towards more negative membrane voltages. In most cases, the maximum conductance, *G*_MAX_, is also increased (Fig. 1D). The *G*(*V*) shift and the *G*_MAX_ increase can be caused by a common site and mechanism of action, but, at least for polyunsaturated fatty acids, which share some functional properties with the resin acids (Ottosson et al., 2014), it has been suggested that these two effects on the cardiac K_V_7.1 channel are mediated via two different parts of the channel (Liin et al., 2018b).

**FIGURE 1.**
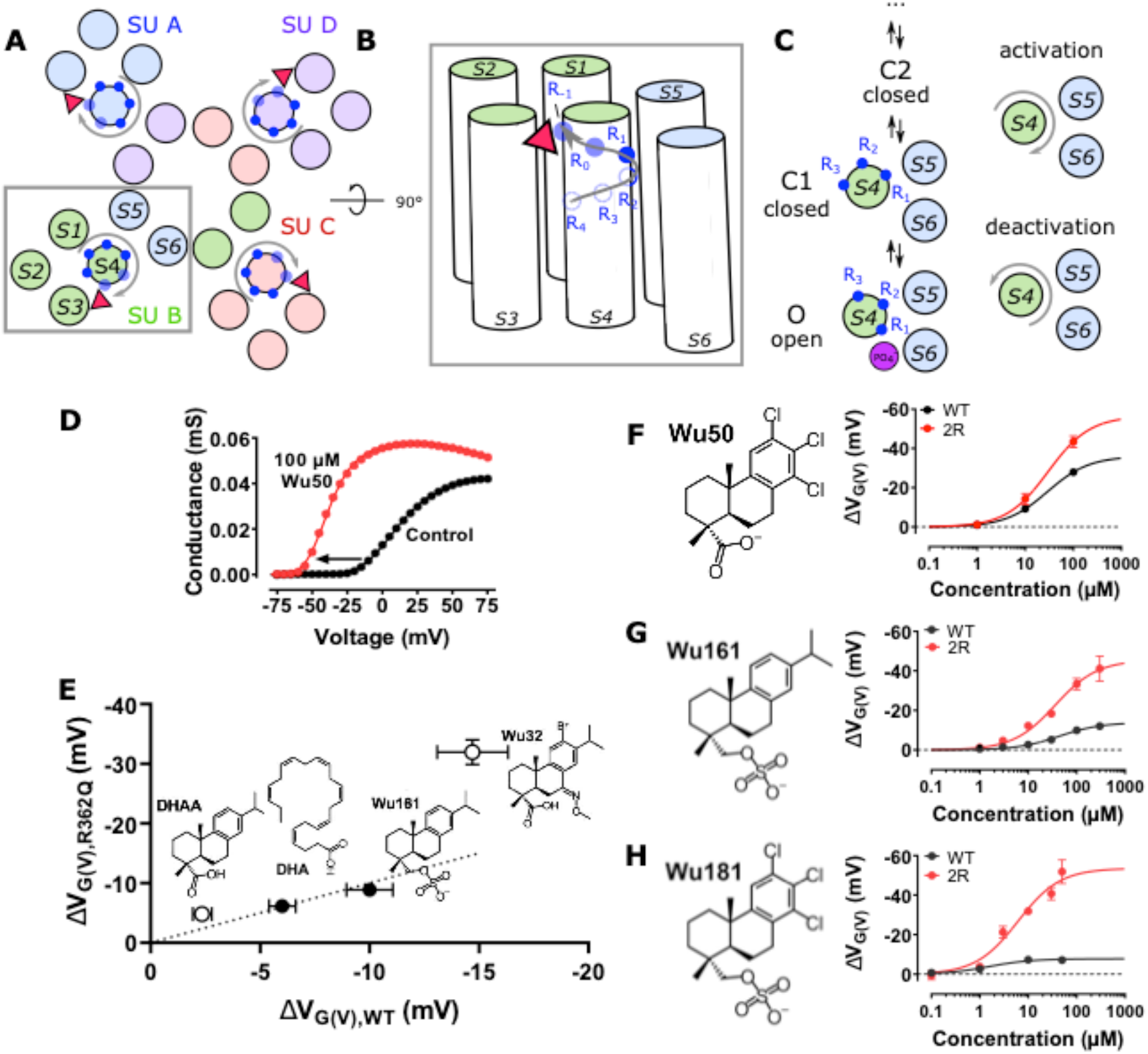
The role of S4 mutations in the Shaker K_V_ channel for the effect of some resin-acid derivatives. **A)** Top and **B)** side view of the S4 helical-screw motion (grey arrow) during VSD activation. Endogenous gating charge arginines denoted by solid blue circles (R362 (=R_1_), R365 (=R_2_), R368 (=R_3_), and R371 (=R_4_)). Additional charges in the 2R motif denoted by transparent blue circles (M356R (=R_-1_), A359R (=R_0_)). Binding of a negatively charged compound (red triangle) in the cleft between S3 and S4 (the S3/S4 site) is hypothesized to favor S4 activation through an electrostatic effect that stabilizes the S4 activated state. The homotetrameric channel assembly is shown in A, with each subunit represented in a different color. A single subunit is shown in B (the pore domain and the VSD are from two different subunits). **C)** Activation of Shaker channels proceeds in several activation steps, between Closed CX and Open O states, during which the S4 helix moves in a ratchet-like upwards and rotative movement (grey arrow). **D)** 100 µM Wu50, at pH 9.0, shifted the *G*(*V*) of the Shaker K_V_ channel with the 2R motif by −40.7 mV. **E)** *G*(*V*) shifts for R362Q vs. WT Shaker K_V_ channels for four compounds. Solid symbols represent compounds for which the shift is similar in the WT and in the R362Q mutant, while empty symbols represent compounds for which the shift is larger in the mutant. The dotted line marks an equal shift in WT and R362Q. **F-H)** Concentration-response curves (Eq. 3). Black, WT Shaker K_V_. Red, Shaker K_V_ with the 2R motif **F)** Wu50, pH = 9.0. WT: *EC*_50_ = 29.1 ± 3.5 µM, Δ*V*_MAX_ = −36.1 ± 1.3 mV, *n* = 3-5. 2R motif: *EC*_50_ = 30.0 ± 5.0 µM, Δ*V*_MAX_ = −56.5 ± 2.6 mV, *n* = 4-10. **G)** Wu161, pH = 7.4. WT: *EC*_50_ = 43.7 ± 6.3 µM, Δ*V*_MAX_ = −13.9 ± 0.6 mV, *n* = 3-4. 2R motif: *EC*_50_ = 36.4 ± 9.0 µM, Δ*V*_MAX_ = −45.5 ± 3.5 mV, *n* = 3-6. **H)** Wu181, pH = 7.4. WT: *EC*_50_ = 1.6 ± 0.6 µM, Δ*V*_MAX_ = −7.8 ± 0.5 mV, *n* = 2-3. 2R motif: *EC*_50_ = 6.1 ± 1.7 µM, Δ*V*_MAX_ = −53.6 ± 4.0 mV, *n* = 4-5. All data Mean ± SEM.

Two experimental findings support the electrostatic channel-opening mechanism: (1) The addition of two positively charged residues at the top of S4 of the Shaker K_V_ channel [M356R/A359R (=R_−1_ and R_0_ in Fig. 1B), from hereon called the 2R motif] increases the channel-opening effect of resin acids (*G(V)*-shift towards more negative membrane voltages; Ottosson et al., 2014, 2015, 2017; Silverå Ejneby et al., 2018). The mechanism we propose involves an increased binding of the resin-acid derivative in the activated/open state due to the proximity of positively charged residues R_−1_ and R_0_ (transparent circles in Fig. 1A,B). (2) Substituting the negative charge on the resin acid by a positive one promotes channel closing (*G(V)*-shift towards more positive membrane voltages; Ottosson et al., 2017), presumably through electrostatic repulsion between the positive charges of the resin acid and of the gating charges in the activated/open state of the channel. Mutagenesis and molecular modeling experiments are consistent with resin-acid derivatives binding between the extracellular portions of S3 and S4, located in the periphery of the VSD (Fig. 1A,B) (Ottosson et al., 2017).

However, some phenomena cannot easily be explained by this simple mechanism: neutralizing the outermost endogenous gating charge R362 to glutamine (R362Q = R1Q) of the Shaker K_V_ channel has surprising effects on the *G(V)-*shifting effects of some resin-acid derivatives (Ottosson et al., 2017; Silverå Ejneby et al., 2018). Using the simple model presented above, we would expect neutralization of R362 to lead to a decreased binding of the compound in the activated state (due to the loss of a positively charged residue close to the binding site) and to thus systematically *decrease* the *G(V)*-shifting effect. Instead, the neutralization of R362 led to an *increase* in effect for two resin acids (dehydroabietic acid (DHAA) and Wu32) and to *no noticeable change* for the more flexible Wu161, and for the polyunsaturated fatty acid docosahexaenoic acid (DHA; a charged lipophilic compound also interacting electrostatically with S4) (Börjesson et al., 2010; Börjesson and Elinder, 2011; Yazdi et al., 2016) (Fig. 1E). Thus, a single binding site, common for all resin-acid derivatives, located at the S3/S4 interface cannot easily explain all the experimental observations.

To be able to design and develop more potent and selective compounds it is imperative to understand the molecular action on ion channels in detail. Therefore, in this work, we sought to explore the molecular mechanism of action of resin-acid derivatives binding with respect to the following questions: (1) Why do resin-acid derivatives have a larger *G*(*V*)-shifting effect on the Shaker K_V_ channel with the 2R motif compared to the WT Shaker K_V_ channel? (2) Why does the R362Q mutation increase the shift for some compounds (e.g. Wu50, Wu32, DHAA) instead of reducing it? (3) Why is the effect of compounds with a longer and flexible stalk (e.g. Wu161, DHA) less affected by the R362Q mutation than the more compact ones?

We performed docking and molecular dynamics simulations (MD) to identify new possible binding sites in the Shaker K_V_ channel and characterize the molecular determinants of binding, and site-directed mutagenesis in the Shaker K_V_ channel with removed fast N-type inactivation (this channel will be referred to wild-type, WT) to test these predictions. We conclude that three possible interaction sites are available around S4 of the Shaker K_V_ channel, making for a complex, state-dependent interaction pattern.

## METHOD

### Expression of Shaker K_V_ channels in *Xenopus laevis* oocytes

The Shaker H4 channel (Accession no. NM_167595.3) (Kamb et al., 1987), with removed N-type inactivation due to a Δ6-46 deletion (ShH4IR, in Bluescript II KS[+] plasmid) (Hoshi et al., 1990), is referred to as the WT Shaker K_V_ channel. Addition of two positively charged arginines (M356R/A359R) at the extracellular top of the voltage sensor increases the effect of charged lipophilic compounds (Ottosson et al., 2014, 2015, 2017). Since the endogenous arginine R362 (=R1) was found to be important for large effect of polyunsaturated fatty acids on the M356R/A359R channel (Ottosson et al., 2014), this channel was referred to as the 3R Shaker K_V_ channel. However, for the resin-acid derivatives studied in the present investigation, R362 reduced the effect. Therefore, we specifically refer to the two added arginines just outside S4 (M356R/A359R) as the 2R motif. Mutations were introduced with site-directed mutagenesis and verified with sequencing as described previously (Börjesson et al., 2010). RNA injection and oocyte handling were carried out as before (Börjesson et al., 2010; Ottosson et al., 2015). All animal experiments were approved by the Linköping’s local Animal Care and Use Committee.

### Electrophysiology

Electrophysiological experiments were made 1–6 days after RNA injection. K^+^ currents were measured with the two-electrode voltage-clamp technique (GeneClamp 500B amplifier; Axon Instruments) as described previously (Ottosson et al., 2015; Silverå Ejneby et al., 2018). The amplifier’s leak compensation was used, and the currents were low-pass filtered at 5 kHz. The control solution contained (in mM): 88 NaCl, 1 KCl, 15 HEPES, 0.4 CaCl_2_, 0.8 MgCl_2_ and pH was set with NaOH. All experiments made in room temperature (20–23°C). The holding potential was set to -80 (or −100 mV if the mutant was not fully closed at −80 mV).

### Resin-acid derivatives

Synthesis of Wu50 (Ottosson et al., 2015), Wu161, and Wu181 (Silverå Ejneby et al., 2018) and has been described previously. Stock solutions were stored at −20°C, and diluted in control solution just prior to experiments. The test solution was added to the oocyte bath manually with a syringe. Wu161 and Wu181 are permanently negatively charged, while the apparent p*K*_a_ in lipid membrane for Wu50 is around 6.5 (Ottosson et al., 2015). To make sure all Wu50 molecules were negatively charged the pH of the control solution was set to 9.0 before the experiments. This pH change had only small effects on channel kinetics and the *G*(*V*) relation. For Wu161 and Wu181 the pH was set 7.4

### Analysis of electrophysiological measurements

The conductance, *G(V)*, was calculated as,

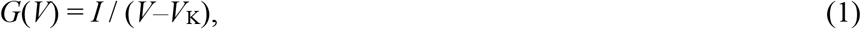

where *I* is the steady-state current measured at the end of each test-voltage sweep (100 ms after onset, Clampfit 10.5, Molecular Devices), *V* the absolute membrane voltage and *V*_K_ the reversal potential for K^+^ ions (set to −80 mV). The conductance was fitted with a Boltzmann equation (GraphPad Prism 5, GraphPad Software, inc),

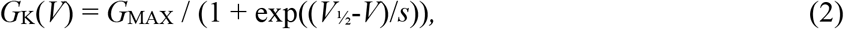

where *G*_MAX_ is the amplitude (maximal conductance) of the curve, *V* the absolute membrane voltage, *V*_½_ the midpoint, *s* the slope. The resin-acid induced *G(V)* shift was calculated as *V*_½_ (compound) − *V*_½_ (control). The relative change in *G*_MAX_ was calculated as *G*_MAX_ (compound)/ *G*_MAX_ (control). The concentration dependence for the *G(V)* shifts were calculated as,

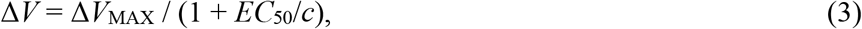

where Δ*V* is the voltage shift, Δ*V*_MAX_ the voltage shift at saturating concentration, *c* the concentration of the tested compound, and *EC*_50_ the concentration at which half maximum response occurs.

### Molecular docking

A previously published model of the Shaker K_V_ channel in the activated/open state was used as the receptor in docking experiments (Yazdi et al., 2016). Initial coordinates for bound resin-acid derivatives Wu50 and Wu161 were obtained using molecular docking to three different binding sites in the open state localized around the outer arginine residues on S4. The docking box was placed around the center of the upper portion of the cleft between S3 and S4 for the S3/S4 site, around the geometric center of the group defined by residues R362, R365, and W454 for the S4/pore site, and around the center of the upper portion of the 4-helix VSD bundle for the top-VSD site. Docking was performed independently to each subunit using Autodock Vina (Trott and Olson, 2010). Since the Autodock Vina scoring function does not account for lipid bilayer environment, the ranking of the docking positions is only approximate. We thus selected docking positions based on a few physical principles: the charged headgroup of the resin-acid derivatives had to localize in the lipid headgroup region, and in direction of positively charged protein groups, while the hydrophobic body localized in the lipid tail region. The coordinates of the ligands docking pose that were subsequently used as initial conditions for the MD simulations were thus different in each subunit and are provided on the Open Science Framework repository (https://osf.io/fw8h9/).

### Molecular dynamics (MD) simulations

The MD simulation system, mimicking the ion channel in its environment, was constructed using the CHARMM GUI Membrane Builder (Wu et al., 2014b). The open state channel and its ligand were placed in a phosphatidylcholine (POPC) bilayer, and the system was hydrated with a 150 mM KCl solution. Mutations (as described in Table 1) were also introduced using CHARMM GUI. The CHARMM36 force field was used to describe interactions between protein (Best et al., 2012;), lipid (Klauda et al., 2010) and ion (Beglov and Roux, 1994) atoms, the TIP3P model was used to describe the water molecules (Jorgensen et al., 1983). The ligand was considered in its charged form, and topology and parameters were prepared using the SwissParam software and general CHARMM force field (CGENFF) (Zoete et al., 2011; Vanommeslaeghe et al. 2010). MD simulations were performed using Gromacs version 5.1.2 (Abraham et al., 2015) on the Beskow supercomputer located at the PDC computer center, Royal Institute of technology KTH. The simulations were performed in sequential steps of minimization, equilibration and production, keeping the default CHARMM GUI parameters (Lee et al., 2016). Initially, 500 ns of simulations were performed for each system. They were then extended for selected systems (Table 1).

**Table 1:** Systems simulated and length of trajectories

| # | Site | Compound | Mutation | Length [ns] |
| --- | --- | --- | --- | --- |
| 1 | S3/S4 | Wu50 | WT | 1000 |
| 2 | S3/S4 | Wu50 | 2R | 500 |
| 3 | S4/pore | Wu50 | WT | 500 |
| 4 | S4/pore | Wu50 | W454A | 500 |
| 5 | S4/pore | Wu50 | R362Q | 500 |
| 6 | S4/pore | Wu50 | W454A/R362Q | 1000 |
| 7 | S4/pore | Wu50 | R362Q/R365Q | 1000 |
| 8 | S4/pore | Wu50 | R362Q/R365Q/W454A | 640 |
| 9 | S3/S4 | Wu161 | WT | 500 |
| 10 | S4/pore | Wu161 | WT | 500 |
| 11 | Top-VSD | Wu50 | WT | 500 |

### MD simulation analysis

Simulation data analysis was performed by scripts using the Python library MDTraj (McGibbon et al., 2015) and are available for download on OSF (https://osf.io/fw8h9/). First, the shortest distance between any heavy atom of the residue in focus and any heavy atom of the ligand were calculated over time. Then the contact frequency was extracted as the fraction of production simulation time spent with any ligand heavy atom within a 4.5 å cutoff of any residue heavy atom. Data is reported for the four subunits independently, and can be seen as four simulation replicates informing on the replicability of the results. Visualization and figure rendering were performed using Visual Molecular Dynamics (VMD) (Humphrey et al., 1996).

### Statistics

Average values are expressed as mean ± standard error of mean (SEM). When comparing two resin acid-induced *G(V)* shifts or EC_50_ values, a two-tailed unpaired t-test was used.

## RESULTS

### The 2R motif increased the maximum *G*(*V*) shift of three resin-acid derivatives

In previous work we suggested that resin-acid derivatives open the Shaker K_V_ channel by binding to the S3/S4 cleft (Fig. 1A,B; Ottosson et al., 2017). One of the arguments supporting this interaction site was that the double-arginine mutation M356R/A359R in the top of S4 (the 2R motif of the Shaker K_V_ channel) increased the *G*(*V*)-shifting effect of resin acids and polyunsaturated fatty acids substantially. The mechanism we proposed for this increased shift presumably involves a direct interaction of M356R and A359R with the compound in the activated/open state, and thus a stabilization of this state relative to the resting/intermediate closed ones (Ottosson et al., 2014, Fig. 1 A,B). To get information on how the apparent affinity (*EC*_50_) and efficacy (maximum *G*(*V*) shift, Δ*V*_MAX_) is affected by the double-arginine mutation M356R/A359R we explored the concentration dependence of Wu50 (Fig. 1F) at Ph 9.0 (a pH at which the compound is fully charged; Ottosson et al., 2015), and of the permanently charged compounds Wu161 (Fig. 1G) and Wu181 (Fig. 1H; Silverå Ejneby et al., 2018) at pH 7.4. The 2R motif increased the Wu50-induced maximum *G(V)* shift by 57% (from −36.1 ± 1.3 mV, *n* = 3-5, to −56.5 ± 2.6 mV, *n* = 4-10), but with no effect in apparent affinity (*EC*_50_(WT) = 29.1 ± 3.5 µM, n = 3-5; *EC*_50_(2R) = 30.0 ± 5.0 µM, *n* = 4-10; Fig. 1F). In contrast to the relatively small increase in *G*(*V*) shift for Wu50, the double-arginine mutation increased the maximum *G*(*V*) shift for Wu161 by 230% (from −13.9 ± 0.6 mV, *n* = 3-4, to −45.5 ± 3.5 mV, *n* = 3-6), and for Wu181 by 590% (from −7.8 ± 0.5 mV, *n* = 2-3, to − 53.6 ± 4.0 mV, *n* = 4-5). While the apparent affinity was not affected for Wu161 (*EC*_50_(WT) = 43.7 ± 6.3 µM, *n* = 3-4; *EC*_50_(2R) = 36.4 ± 9.0 µM, *n* = 3-9), it seemed to slightly decrease for Wu181, but this effect was not statistically significant (*EC*_50_(WT) = 1.6 ± 0.6 µM, *n* = 2-3; *EC*_50_(2R) = 6.1 ± 1.7 µM, *n* = 4-5). In summary, while the maximum *G*(*V*) shift was about equal for all three compounds on the Shaker K_V_ channel with the 2R motif (−46 to −57 mV), the maximum *G*(*V*) shift on the WT Shaker K_V_ channel was much smaller for the two flexible stalk compounds (Wu161 and Wu181; −8 and −14 mV) compared with the more compact Wu50 (−36 mV). The apparent affinity, however, was not dependent on channel mutations, but compound-dependent, with Wu181 being the most potent compound. Because the oocytes in many experiments did not tolerate higher concentrations of Wu181, we used Wu50 and Wu161 for the remainder of this study.

### The 2R motif increased the binding of Wu50 to the S3/S4 cleft

To gain molecular level insights into the effect of the 2R mutation, we turned to molecular docking and MD simulations. While MD simulations do not assess compound efficacy, they provide an indication of the kinetics of unbinding from the active state of the Shaker K_V_ channel through the observation of a few stochastic events. Since the channel considered here is a homotetramer, each simulation initiated with a slightly different docking pose in each subunit provides four quasi-independent observations, and each subunit can be considered as a replicate simulation. Overall, if compounds tend to detach easily from their initial binding pose, this may indicate that there is no free energy minimum at the initial docking pose location. When compounds seem to stay stably bound, this can serve as an indication of a high free energy barrier towards unbinding, and thus indirectly of a putative high binding affinity. We also note that docking and MD simulations are limited to the only experimentally available channel state, the open O state. Thus, the conclusions drawn are not based on explicit observations made in other states. This is unfortunate since our model relies on the relative affinity of compounds to open versus closed states. Yet, by combining insights from the computational and mutagenesis work, we are able to gradually build a mechanistic model of the effect of the resin acid compounds on our model channel.

Docking of Wu50 to the S3/S4 site of the fully activated/open WT Shaker K_V_ channel, followed by a 1-µs long simulation of this system, revealed a pose in which the negatively charged carboxyl group of Wu50 was localized in the headgroup region of the lipid bilayer and interacted with polar residues Thr326 or Thr329 (in the extracellular end of S3), or with the positively charged residue Lys266 (in the S1-S2 loop), while its hydrophobic body partitioned in the lipid tail region and was in contact with hydrophobic residues of the S3/S4 upper cleft Ile325, Ala359, Ile360 and Ile364 (Fig. 2A-D). The binding pose, however, appeared quite unstable in three out of four subunits (Fig. 2B). In one of the subunits, the orientation of the compound even changed drastically, assuming a position parallel to the membrane surface, with the negatively charged headgroup reaching out of the pocket towards R362 (Subunit C, Fig. 2A,C). In two subunits (Subunits A and B, Fig. 2A-C), the compound eventually detached from the binding site after a few hundreds of nanoseconds. We noted that this occurred in the two subunits where Lys266 pointed away from the binding site, while in subunit D, where Lys266 pointed towards the binding site, the binding of Wu50 remained stable over the entire length of the MD simulation. We thus conclude that binding to the S3/S4 cleft appeared relatively weak and that the charged residues present at the top of the VSD played a role in keeping Wu50 close to the channel.

**FIGURE 2.**
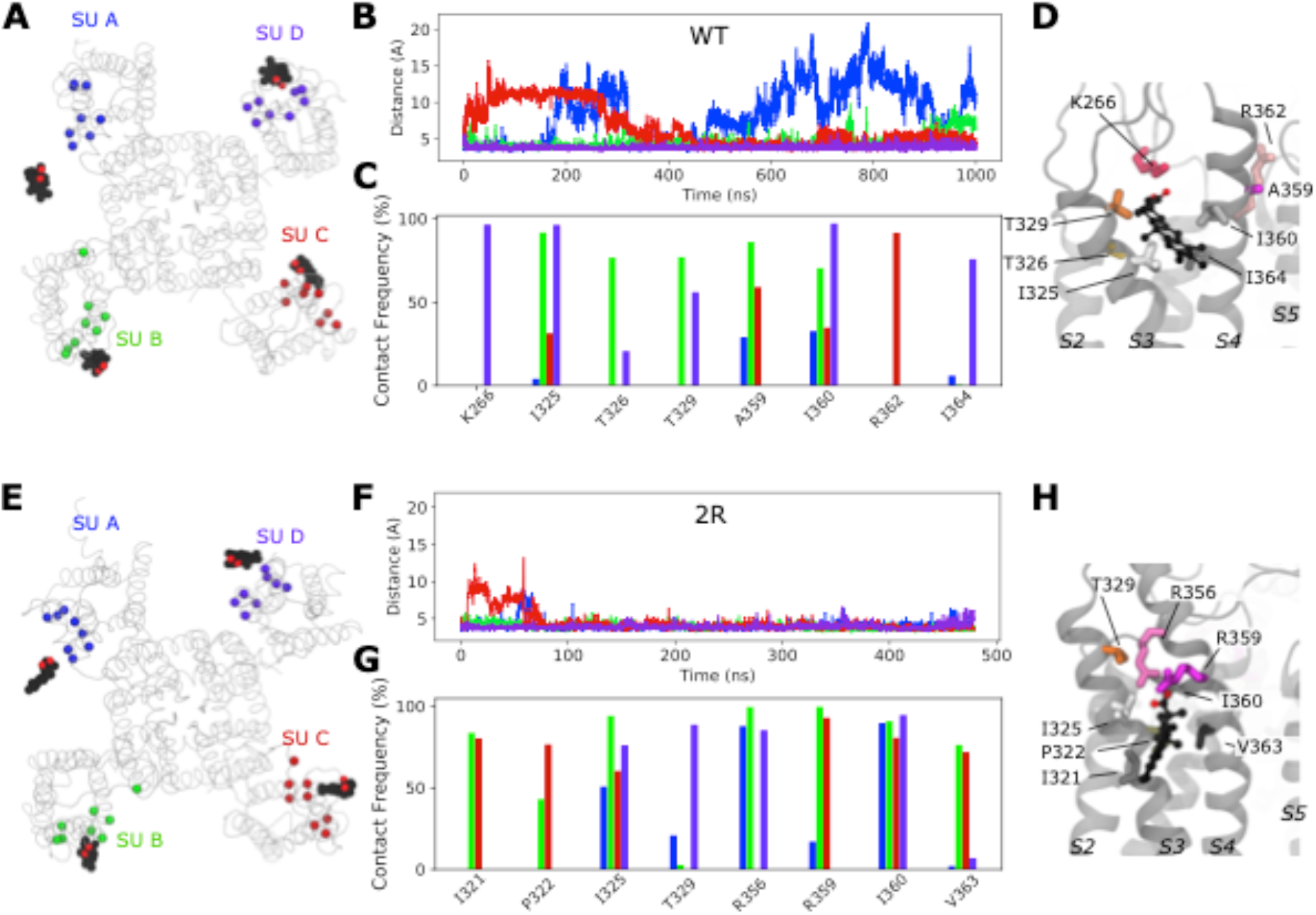
Molecular insight into binding of Wu50 to the S3/S4 site. **A)** Top view of a representative snapshot of the interaction between Wu50 and the S3/S4 site in the WT Shaker K_V_ channel. The channel is shown as grey ribbons, the Wu50 compounds are displayed using a space-filling representation, each atom type colored differently (black: C or Cl; red: O) Residues’ C_α_coming in contact with Wu50 in at least one of the channel’s subunits during MD simulations are represented as spheres and colored according to the subunit they belong to. **B)** Distance between closest heavy atoms of Wu50 and ILE360 along a 1-µs long MD simulation of the WT Shaker K_V_ channel. Each of four subunits is depicted in a different color, following the color scheme presented in panel A. **C)** Contact frequency between any heavy atom of Wu50 and S3/S4 site residues in the WT Shaker K_V_ channel MD simulation. Each of the four subunits is depicted in a different color, following the color scheme presented in panel A. For each residue, each bar corresponds to the contact frequency in one of the four subunits. **D)** Zoomed-in side view of the S3/S4 site for the WT Shaker K_V_ channel. The residues identified in the contact frequency analysis are shown as sticks. Apolar, polar, and positively charged residues are represented as sticks and depicted in shades of grey, orange, and pink, respectively. Wu50 compounds are displayed using a CPK representation. **E)** Top view of a representative snapshot of the interaction between Wu50 and the S3/S4 site in the Shaker channel with the 2R motif. **F)** Distance between closest heavy atoms of Wu50 and ILE360 along a 500-ns long MD simulation of the 2R motif-channel system. **G)** Contact frequency between any heavy atom of Wu50 and S3/S4 site residues in the 2R motif channel simulation. **H)** Zoomed-in side view of the S3/S4 site for the channel with the 2R motif. Colors and representations in E-H are the same as in A-D.

To further test the role of positively charged residues on S4, we docked Wu50 to the S3/S4 cleft in the fully activated/open Shaker K_V_ channel containing the 2R motif (Fig. 2E-H). In all subunits, the negative charge of Wu50 quickly oriented towards M356R and/or A359R (Fig. 2G,H). This interaction maintained the hydrophobic body in contact with I360 for more than 80% of the simulation time and with I325 for more than 50% of the simulation time in all subunits (Fig. 2G). While different subunits displayed different behaviors, this site clearly appeared more stable than in the WT Shaker K_V_ channel. The electrostatic potential at the pocket, which is heavily influenced by the presence or absence of positively charged residues, thus seems to control the affinity of the negatively charged compound for the S3/S4 binding site.

As mentioned above, we were not able to conduct explicit docking and MD simulations in the C1 closed state of the channel. However, as S4 moves down, we expect the charges to leave the binding site both in the WT and the 2R channel, and the affinity for this site to be weak in this state. The difference in binding affinity for the two states in thus larger for the 2R channel than for the WT and thus explains why the *G(V)* shifting effect is larger for the mutant.

Electrophysiology data (Fig. 1F) indicate that the apparent efficacy of Wu50 towards the Shaker K_V_ channel with or without the 2R motif differs while its affinity to both channels is similar. The apparent difference in affinity for the activated/open state inferred from the MD simulations, as well as the unexplained behavior of the R362Q mutant for some resin acids (Fig. 1E), could be an indication of the presence of other binding sites in the channel’s periphery.

### A S4/pore pocket is important for Wu50 effects

In the active state (in the absence of a modulator compound), R362 (and R365) of the Shaker K_V_ channel interacts electrostatically with the PO_4−_ group of zwitterionic POPC lipids forming the membrane (Fig. 3A) (Kasimova et al., 2014). This interaction disappears when the channel deactivates, that is as the S4 helix moves down into the intermediate closed states in a helical-screw motion (Tombola et al., 2007). Indeed, during the first steps of deactivation, R362 and R365 leave the lipid headgroups to interact with negatively charged amino acids located on helices S1-S3 (Delemotte et al., 2011; Henrion et al., 2012). It follows therefrom that when the upper leaflet contains negatively charged lipids, the activated state is stabilized through an electrostatic effect, which is reflected through a *G(V)*-shift towards more negative membrane voltages. When lipid headgroups are absent (as is the case when the membrane is made of ceramide lipids, for example), on the other hand, the activated state is destabilized due to the lack of binding site for R362 and R365 in the activated state (Kasimova et al., 2014). The environment around the fully activated S4 helix, close to R362 and R365, thus seems to be able to accommodate compounds that are both hydrophobic and negatively charged.

**FIGURE 3.**
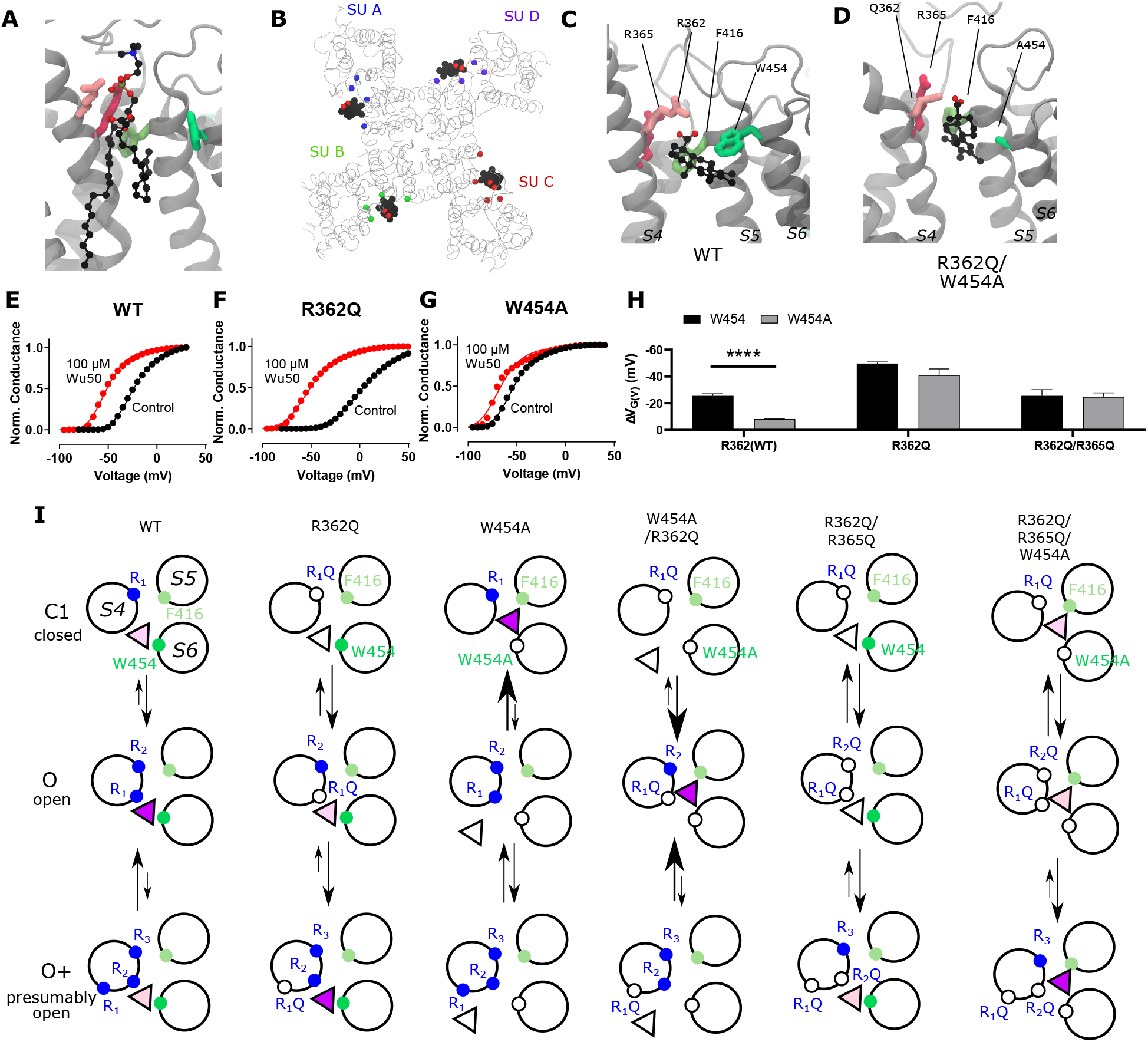
A site located at the S4/pore interface is important for the effect of Wu50 on the WT Shaker K_V_ channel. **A)** Side view of the S4/pore site for the WT Shaker K_V_ channel in the absence of resin-acid derivative. A POPC molecule is occupying the binding pocket. POPC is displayed using a CPK representation, each atom type colored differently (black: C; red: O; blue: N; brown: P; hydrogens are omitted for clarity). Important interacting residues are represented as sticks (shades of green: aromatic residues, shades of pink: positively charged residues) **B)** Top view of a representative snapshot of the interaction between Wu50 and the S4/pore site in the WT Shaker K_V_ channel. The channel is shown as grey ribbons, the Wu50 compounds are displayed using a space-filling representation, each atom type colored differently (black: C or Cl; red: O) Residues’ C_α_ coming in contact with Wu50 in at least one of the channel’s subunits during MD simulations are represented as spheres and colored according to the subunit they belong to. **C)** Zoomed-in side view of the S4/pore site for the WT Shaker K_V_ channel in the presence of the resin-acid derivative Wu50. Wu50 is displayed using a CPK representation, each atom type colored differently (black: C; red: O). The rest of the representation is the same as in A. **D)** Zoomed-in side view of the deeper S4/pore site for the W454A/R362Q mutant channel in the presence of the resin-acid derivative Wu50. **E-G)** Representative normalized *G*(*V*) curves for Wu50-induced effects on different Shaker K_V_-channel mutants. 100 µM, pH = 9. **H)** Wu50-induced *G*(*V*) shifts for different Shaker K_V_-channel mutants. 100 µM, pH = 9. Mean ± SEM (*n* = 3–6). **** = p < 0.0001. **I)** Scheme of putative state-dependent interactions between Wu50 in the S4/pore site in the WT and the various mutants investigated. The endogenous gating-charge arginines (R362 (=R_1_), R365 (=R_2_)) are denoted by filled blue circles, F416 and W454 by filled green circles. Mutations are denoted by empty circles. Mutation of W454 to Ala opens a deeper binding site in the vicinity of F416. Putative binding affinities for the site range from weak (white triangles) to medium (light purple triangle) to strong (dark purple triangle). Transition between states are represented by arrows. Relative stabilization of a state through binding of the compound to this site leads to increased transitions to this state. Stabilization by medium binding increases the transition slightly (medium size arrow) and stabilization by strong binding increases the transition greatly (large size arrow). A reduction in transition probability is marked by a smaller arrow. The overall *G*(*V*) shift is due to the stabilization of Open states (O and O+) relative to closed states (C1 and other closed states not represented here).

We thus hypothesized that the S4/pore pocket may harbor a potential stable binding site for a negatively charged compound such as Wu50, which has a large effect on the WT Shaker K_V_ channel (Fig. 1F) but appears to interact weakly with the activated channel’s S3/S4 cleft in absence of the 2R motif (Fig. 2B). To test this hypothesis, we docked Wu50 to the fully activated/open WT Shaker K_V_ channel, centering the docking box close to the PO_4−_ group coordinating R362. The 500 ns MD simulations initiated from these poses indicated that Wu50 tended to stay in this pocket (Fig. 3B,C, Supplementary Fig. S1, S2). The negatively charged carboxyl group of Wu50 made a close interaction with the positively charged R362. In addition, Wu50 displayed a prominent interaction with a tryptophan residue located in the extracellular end of S6 (W454) (Fig. 3C, Supplementary Fig. S1, S2). In some subunits, the flat aromatic sidechain ring of W454 formed a tight parallel interaction with the flat tri-chlorinated aromatic skeleton ring of Wu50 (Fig. 3C). A phenylalanine residue located in the extracellular end of S5 (F416) provided an interaction on the other side of the binding site (Fig. 3C, Supplementary Fig. S1, S2). The stable and simultaneous interaction with R362 and W454 would be lost as the channel deactivates and S4 moves down while R362 rotates away from the S4/S5 cleft. We thus hypothesize that binding to this site, which we call the S4/pore site, stabilizes the VSD activated state and therefore promotes channel opening.

To experimentally test a potential binding site between R362 and W454 we mutated these residues one by one and in combination. As mentioned in the introduction, removing the top gating charge (R362Q) did not decrease the effect but increased it (Fig. 3F, H, Supplementary Table S1), from −25.5 ± 1.4 mV (*n* = 5) to −49.6 ± 1.2 mV (*n* = 3). In contrast, if the tryptophan 454 in S6 was mutated to an alanine, the *G(V)*-shift was substantially reduced (Fig. 3G, H, Supplementary Table S1), from −25.5 ± 1.4 mV (*n* = 5) to −8.1 ± 0.4 mV (*n* = 5). Simultaneously mutating W454 and R362 (R362Q/W454A) increased the effect compared to WT (Fig. 3G, Supplementary Table S1, −41.0 ± 4.4, *n* = 5).

To explain these observations we put forward two possible mechanisms which are not mutually exclusive and may coexist: (1) The mutation *per se* can increase the *G*(*V*) shifting effect of a compound without affecting the interaction between the compound and the channel. One example of this is the ILT mutation (Smith-Maxwell et al. 1998) which increases the *G*(*V*) shift induced by polyunsaturated fatty acids and resin acids through a separation of early and late voltage-dependent steps in the channel-activating sequence (Börjesson & Elinder, 2011; Ottosson et al., 2017). As described in Supplementary Information, however, the effects of the R362Q mutation on the separation of early and late voltage-dependent steps are mild compared with the ILT mutant and the estimated effect of this mechanism on the *G*(*V*) shift appears rather small. (2) The other explanation is that S4 can rotate further into a hyper-activated O+ state, revealing a putative binding site between R365 and W454 (Fig. 3I). This site is suggested to be revealed by the R362Q mutation and is otherwise unstable (Fig. 3I, column 2).

Altogether, according to the model mentioned previously, it is the selective binding of the compounds to the O and O+ state relative to the C1 and other closed states that determine the magnitude of the *G*(*V*) shift. We detail here how this model can be used to interpret the *G*(*V*) shifts in the different mutants (Fig. 3I): in the WT Shaker K_V_ channel, binding of Wu50 is strong in the O state and weaker in the C1 and O+ states (first column in Fig. 3I). Binding to C1 is presumably weak since R_1_ is located at a distance from the binding site. Binding to O+ is also presumably weak, but due to other reasons possibly linked to the presence of R_1_ further outwards. In contrast, the R362Q mutation reduces binding to the C1 and O states (due to the loss of a positive charge in the binding site) while increasing binding to the O+ state (second column in Fig. 3I, Supplementary Fig. S2, S3). This mechanism is consistent with our previous results on two other resin acids (DHAA and Wu32) showing an increased *G*(*V*) shift upon mutation of R362 (Ottosson et al., 2017; Silverå Ejneby et al., 2018, Fig. 1D). MD simulations of the O state W454A mutant showed an increase in instability of Wu50 binding at this site relative to WT. We interpret this as a sign of reduced binding due to the loss of interactions with W454 (third column in Fig. 3I; Supplementary Fig. S2, S4). Additionally, MD simulations of the O-state R362Q/W454A mutant showed that Wu50 finds a deeper stable binding site between F416 (which replaced the interaction with aromatic residue W454) and R365 (Fig. 3D, column four in Fig. 3I, Supplementary Fig. S2, S5). To explain the increased shift in the R362Q/W454A mutant relative to WT, we need to hypothesize that this deep binding site is not occupied in the C1 and O+ state. By symmetry, the site is occupied in the W454A mutant in the C1 state (thanks to the proximity of R_1_) and not the O and O+ state, providing an explanation for the reduced *G*(*V*) shift for this mutant (column three in Fig. 3I).

To experimentally test the idea that F416 is involved in a deep binding site, we explored R362Q/W454A/F416A. Unfortunately, Wu50 reduced *G*_MAX_ by 83%, and we could therefore not reliably measure the G(V) shift (Supplementary Table S1).

To experimentally test the idea that R365 is involved in state-dependent binding, we also neutralized R365 (R362Q/R365Q). The *G(V)* shift was very much reduced from −49.6 ± 1.2 mV (*n* = 3; R362Q) to −25.5 ± 4.6 mV (*n* = 5; R362Q/R365Q) (Fig. 3H). Using our mechanistic model further, we propose that a weak binding in the O+ state (due to electrostatic interaction with R3 at a distance), but not in the O and C1 states, could explain this effect (column five in Fig. 3I). In line with this, the R362Q/R365Q/W454A mutation showed the same sensitivity to Wu50 (−24.7 ± 3.0, *n* = 6; Fig. 3H, Supplementary Table S1) as R362Q/R365Q. Since we know that the W454A mutation leads to binding to the deeper site, by symmetry, we propose that binding is weak in the C1 and O state but strong in the O+ state, resulting in a similar relative stabilization of states in R362Q/R365Q and R362Q/R365Q/W454A (Column six in Fig. 3I, Supplementary Fig. S2, S6, S7).

It should be noted that mutations in the S4/pore pocket also altered the effect of the compounds on *G*_MAX_ (Supplementary Information and Table S1). This will be further explored in a later study.

### The 2R motif rescues the W454A effect

When the 2R motif was combined with the *G(V)*-shift-attenuating mutation W454A, the attenuation was gone (Fig. 4A, C, Supplementary Table S1). This suggests that the 2R motif trumps the effects of a compound bound to the S4/pore pocket. Combining the 2R motif with the R362Q mutation increased the *G*(*V*) shift even more (from −40.0 ± 2.7 mV (*n* = 8), to − 54.0 ± 2.3 mV (*n* = 4; Fig. 4B, C, Supplementary Table S1), and combining the 2R motif with R362Q/W454A mutation increased the G(V) shift from -41.0 ± 4.4 (*n* = 5) to −56.6 ± 5.8 (*n* = 4; Fig. 4C, Supplementary Table S1). Thus, we hypothesize that in a channel with the 2R motif, the S3/S4 site dominates and the S4/pore pocket has no additional effect. However, in the absence of R362 (and the 2R motif), as described above, we hypothesize that the binding of the compound in the S4/pore pocket increases the *G(V)* shift through the recruitment of R365, which has a tendency to pull S4 into the stable O+ state, and cause opening of the pore at lower voltages (Fig. 3F, I). The 2R motif is therefore not *per se* needed for increasing the *G(V)*-shifting effect of resin acids.

**FIGURE 4.**
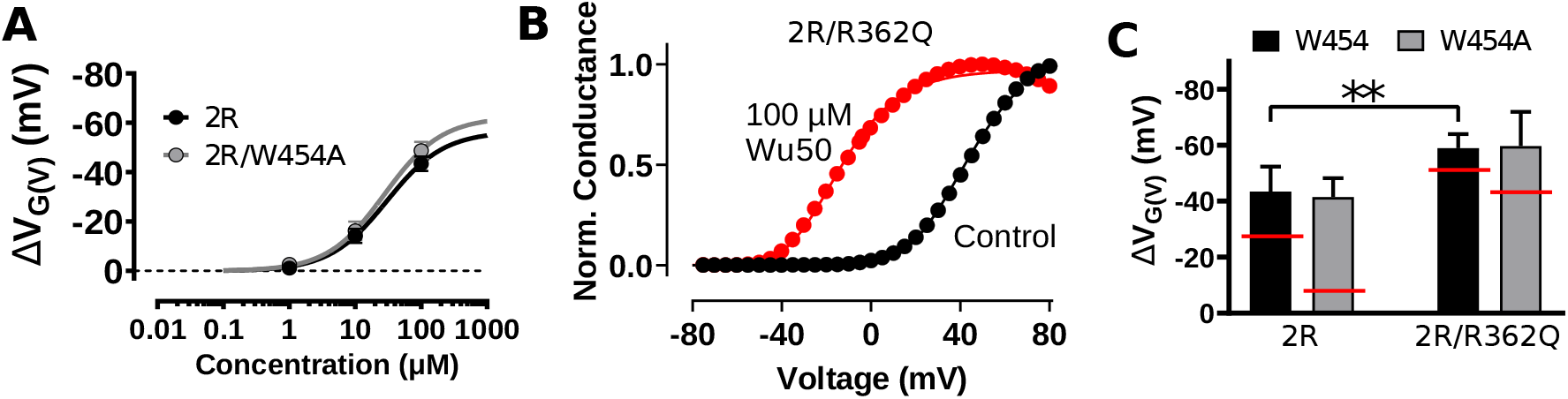
The 2R motif overrides the effect from the S4/pore site. **A)** Concentration-response curves for Wu50 at pH 9.0. 2R motif: *EC*_50_ = 30.0 ± 5.0 µM, Δ*V*_MAX_ = −56.5 ± 2.6 mV, *n* = 4-10; 2R motif/W454A: *EC*_50_ = 27.9 ± 9.2 µM, Δ*V*_MAX_ = −62.3 ± 6.5 mV, *n* = 3. Mean ± SEM. **B)** 100 µM Wu50 at pH 9.0 shifts *G*(*V*) for the 2R motif/R362Q Shaker K_V_ channel results by −59.0 mV. **C)** Wu50-induced *G(V)* shifts for Shaker K_V_ channel mutants as indicated. 100 µM, pH = 9. Mean ± SEM (*n* = 4-10).). **, p < 0.01. The red lines denote the shifts in the absence of the double arginine mutation (M356R/A359R = 2R).

### A longer and more flexible stalk alters Wu161 binding to the S4/pore site

Wu161 and Wu181, which have their negative charge on a three-atom long and slightly flexible stalk, have smaller effects on the WT Shaker K_V_ channel compared with Wu50, but a very large effect when the 2R motif is introduced (Fig. 1F-H). This suggests that the stalk compounds Wu161 and Wu181 exert their main effect on the Shaker K_V_ channel with the 2R motif from to the S3/S4 site.

MD simulations initiated with Wu161 docked to the S3/S4 site of the fully activated/open WT Shaker K_V_ channel did not display a prominently different behavior from Wu50, with two out of four compounds displaying a tendency to leave their binding site (Fig. 5A,B). On the other hand, docking of Wu161 to the S4/pore site showed substantial differences from Wu50 (Fig. 5C,D). The binding to the WT activated/open Shaker K_V_ channel appeared overall less stable: Wu161 left the binding site in one of the subunits and the minimum distance from the compound to R362 appeared generally larger and displayed more fluctuations than when with Wu50. Indeed, scrutinizing the trajectories led to the observation that Wu161 did not fit as tightly in this binding site because its longer stalk made it impossible for their hydrophobic body to interact with W454 via pi-stacking at the same time as their negatively charged headgroup interacted with the positively charged group of R362 (Fig. 5C).

**FIGURE 5.**
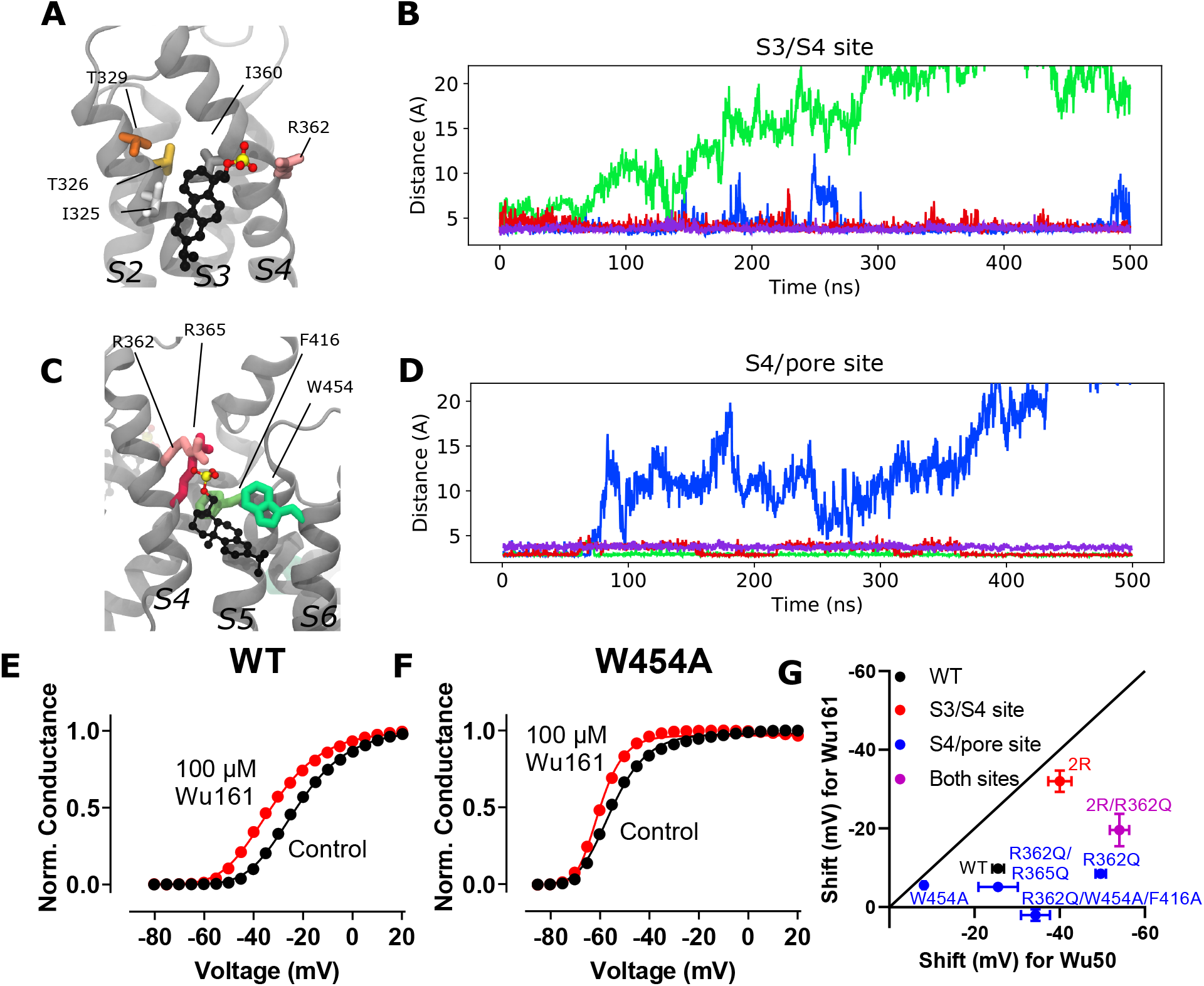
Wu161 binding to the S3/S4 and the S4/pore sites. **A)** Zoomed-in side view of the S3/S4 site in the presence of wu161. The channel is shown as grey ribbons, the Wu161 compounds are displayed in CPK representation, each atom type colored differently (yellow: S; black: C; red: O) Apolar, polar, positively charged and aromatic residues are represented as sticks and depicted in shades of grey, orange, pink, and green, respectively. **B)** Distance between closest heavy atoms of Wu161 and I360 along a 500 ns long MD simulation of the WT Shaker K_V_ channel system. Each of four subunits is depicted in a different color. **C)** Zoomed-in side view of the S4/pore site in the presence of Wu161. Representations are the same as in A. **D)** Distance between closest heavy atoms of Wu161 and R362 along a 500 ns long MD simulation of the WT Shaker K_V_ channel system. Each of four subunits is depicted in a different color. **E-F)** *G*(*V*) curves before (black) and after (red) application of 100 µM Wu161 to the channels. *G*(*V*) shifts = −10.4 mV and −4.7 mV respectively. pH = 7.4. **G)** Correlation between *G*(*V*) shifts for Wu50 and Wu161 (100 µM) on different Shaker mutants. Error bars represent Mean ± SEM (*n* = 4). The solid black line marks an equal *G*(*V*) shift for Wu50 and Wu161.

Mutating W454A reduced the *G(V)* shift for Wu161 (from −9.8 ± 1.1 mV, *n* = 5, to −5.6 ± 0.8 mV, *n* = 4; Fig. 5E-F; Supplementary Table S1), suggesting that Wu161 does bind to the S4/pore site, but its effect on the WT Shaker K_V_ channel is smaller than that of Wu50 (−25.5 ± 1.4; *n* = 5), which may indicate that binding of this compound to this site is less favorable than that of Wu50, in line with the MD simulation results. Removing R362 does not increase the *G(V)* shift (−8.5 ± 0.5, *n* = 3), as it does for Wu50 (−49.6 ± 1.2, *n* = 3; Fig. 5G, Supplementary Table S1). This may indicate that while Wu50 fits well between R365 and F416 in the absence of R362 and causes channel opening, Wu161 either does not bind here, or when bound, cannot keep the channel in the open state. In an attempt to further weaken a possible binding of Wu161 to the S4/pore site we tested the triple mutation R362Q/F416A/W454A. The shift was completely eliminated (Supplementary Table S1). This mutant was, however, surprisingly sensitive to changes in pH (completely blocked at pH 9) and could therefore not be tested for Wu50.

The R362Q/R365Q double mutant reduced the Wu161-induced *G(V)* shift compared with R362Q (from −8.5 ± 0.5, *n* = 3, to −5.2 ± 1.2, *n* = 4; Fig. 5G, Supplementary Table S1). Such a reduced effect may indicate that this double mutation abolishes binding of this compound entirely to the S4/pore site. To conclude, the flexible compound Wu161 seems to bind to the S4/pore site, but the binding is less stable than for the more rigid compound Wu50, and the *G*(*V*) shifting effects are thus substantially smaller (Fig. 5G, blue symbols). In contrast, Wu161 bound to the channel with the 2R motif has a large *G*(*V*) shifting effect (−32.0 ± 2.7 mV, *n* = 6), indicating stable binding to the S3/S4 site when the 2R motif is present (Fig. 5G, red symbol; Supplementary Table S1).

### A possible binding site at the top of the VSD bundle

Scrutinizing further the surroundings of S4, we considered a third putative binding pocket in the center of the extracellular region of the VSD (which we call the top-VSD site, Fig. 6). Indeed, positive charges from S4 (R365, R368, and even R371=R4) are exposed to the extracellular solution, and this site has been shown to be druggable for other classes of compounds (Liin et al., 2018a), and in related channels (Ahuja et al., 2015; Li et al., 2013; Peretz et al., 2010). We thus docked Wu50 to this putative site surrounded by the S1, S2, S3, and S4 helices. Binding there in the O state appears stable in MD simulations (Fig. 6.C), the negative charge of Wu50 being close to R365 and R368. Other residues often in contact with the compound include F280 and E283 in S2, and T326 in S3 (Fig. 6). Experimental tests of the role of these residues via mutagenesis proved difficult; the mutations only showed very small currents or the compound blocked the current, thus precluding studies of the *G*(*V*) shifting effect (see Supplementary Table 1).

**FIGURE 6.**
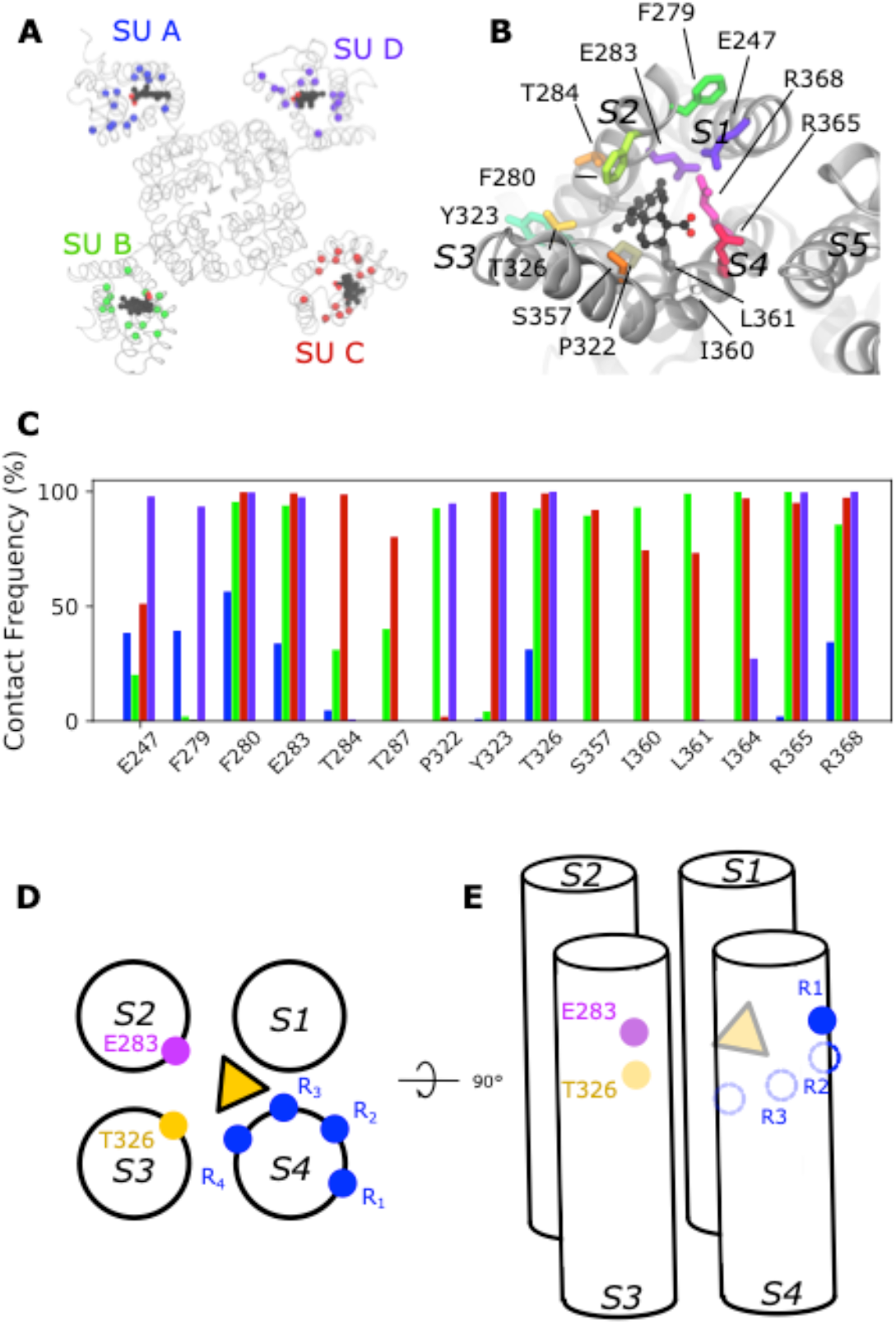
A possible binding site at the extracellular side of the VSD. **A)** Top view of a representative snapshot of the interaction between Wu50 and the top-VSD site in the WT Shaker K_V_ channel. The channel is shown as grey ribbons, the Wu50 compounds are displayed as space-filling, each atom type colored differently (black: C or Cl; red: O) residues are represented as sticks. Residues’ C_α_ coming in contact with Wu50 in at least one of the channel’s subunits during MD simulations are represented as spheres and colored according to the subunit they belong to. **B)** Zoomed-in top view of the top-VSD binding site for the WT Shaker K_V_ channel in the presence of resin-acid derivative Wu50. **C)** Contact frequency between any heavy atom of Wu50 and the top-VSD site residues in the WT Shaker K_V_ channel simulation. For each residue, each bar corresponds to the contact frequency for one of the four subunits, depicted in a different color, following the color scheme presented in Fig. 1. **D)** Top and **E)** side schematic view of the putative effect of resin-acid derivative binding to the top-VSD site (yellow triangle) The endogenous gating charge arginines (R362 (=R_1_), R365 (=R_2_), R368 (=R_3_) and R372 (=R_4_)) are denoted by blue filled circles. The positions of E283, and T326 are denoted by pink and yellow circles, respectively.

## DISCUSSION

The helical-screw motion of S4 combined with the periodic arrangement of arginines every helical turn defines three potential binding sites for resin acids and their derivatives, that can be more or less occupied depending on state- and compound-dependent available interaction motifs (Fig. 7).

**FIGURE 7.**
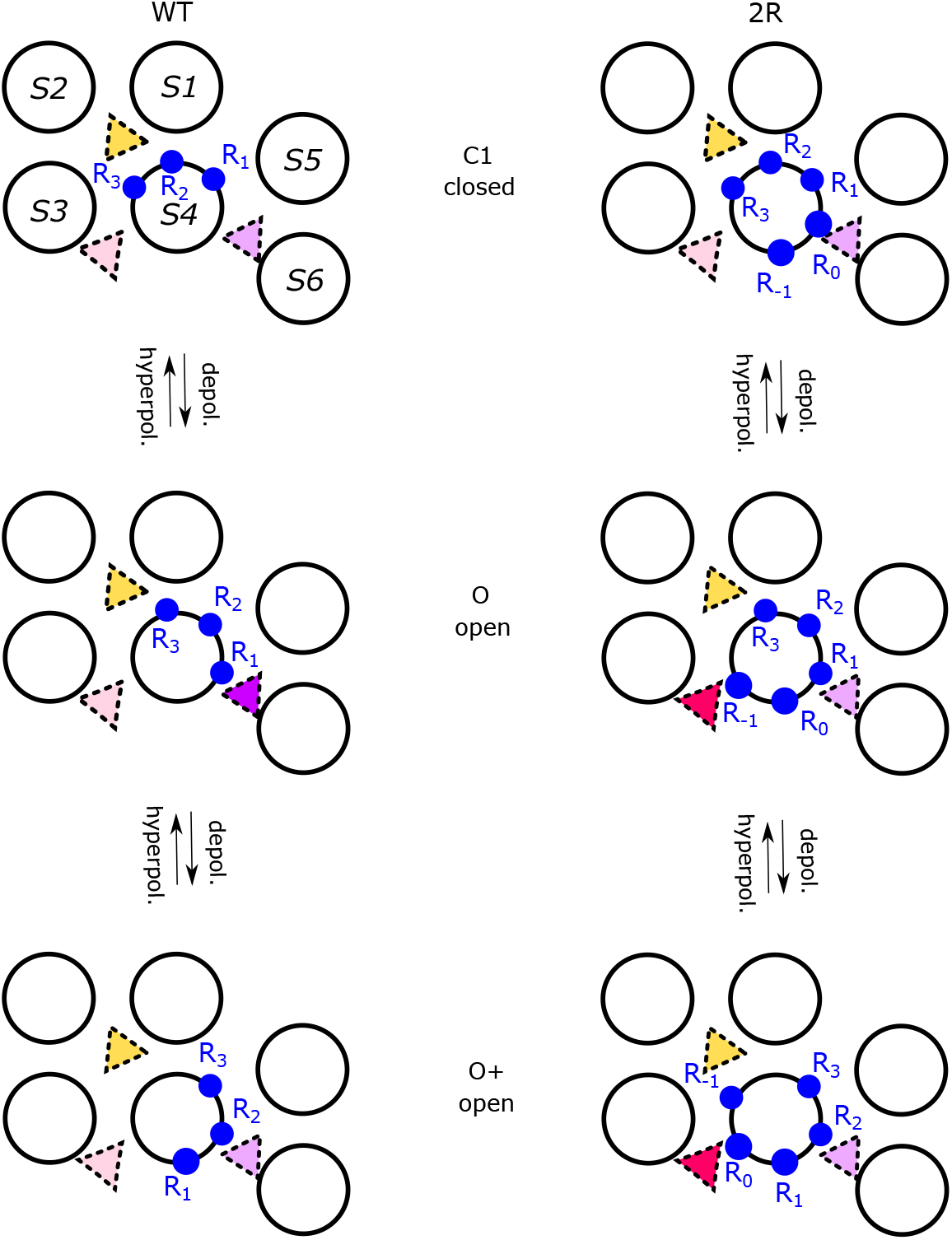
Three putative resin-acid binding sites act on activation via differential stabilization of S4 in various activation states. Top views of the motion of S4 during the last steps of activation occurring in one subunit (N.B. the pore domain and VSD are from two different subunits) in the WT Shaker K_V_ channel (left) and the 2R mutant (right), from the C1 closed state (top row) to the open O state (middle row) to the putative open O+ state (bottom row). Binding to the S3/S4 site is represented as a red triangle, to the S4/pore site as a purple triangle and to the top-VSD site as a yellow triangle. The charged residues are shown as blue filled circles. In the last intermediate (C1) closed state, both for the WT and the 2R channel, binding to the S3/S4 site is weak (light red triangle), binding to the S4/pore site is weak (light purple triangle) and binding to the top-VSD site is strong (yellow triangle). In the open O state, binding to the S3/S4 site is weak (light red triangle) in the WT but strong (bright red triangle) in 2R due to the proximity of R_-1_, binding to the S4/pore site is strong (bright purple triangle) in the WT due to the proximity of R_1_ but presumably weak in the 2R (light purple triangle) and binding to the top-VSD site is strong (yellow triangle) in both the WT and the 2R mutants. In the hyperactivated/open O+ state, binding to the S3/S4 site is weak (light red triangle) in the WT but presumably strong (bright red triangle) in 2R due to the proximity of R_0_, binding to the S4/pore site is weak (light purple triangle) in both the WT and the 2R mutants and binding to the top-VSD site is strong (yellow triangle) in both the WT and the 2R mutants.

### Three tentative binding sites - a summary

Binding of resin-acid compounds to the S3/S4 site (Fig. 7, red triangle) tends to favor S4 activation (Fig. 7, displacement downwards from the C1 to the O state) through an electrostatic effect. Indeed, this site is particularly occupied in the activated O state when the 2R motif is introduced, as evidenced by MD simulations and thanks to interactions between the compound and R_-1_. It would also be favorably occupied in a hypothetical hyper-activated O+ state channel thanks to the recruitment of R_0_ to this site, although whether the affinity for this site in this state is high or medium remains to be investigated. Binding to this site in the absence of the 2R motif in any state, on the other hand, is seemingly relatively unstable: in the activated O state R362 (R_1_) and R365 (R_2_) are located on the opposite side of the S4 helix and R368 (R_3_) and R371 (R_4_) are located too far down, a situation not drastically modified by a transition to the C1 or the O+ states.

We further hypothesize that binding to the S4/pore site (Fig. 7, purple triangle) tends to favor S4 activation (transition from C1 to O state) through direct binding to R_1_. The aromatic ring-like body of the resin-acid derivative anchors to W454, or, in its absence, to F416 (Fig. 3C, D, I). Removing R_1_ can enhance the effect of binding to this site since its absence presumably favors a transition to an even more activated state O+ where R_2_, which is usually buried further into the S4/S5 crevice, is further pulled upwards. What happens in this site in the presence of the 2R motif is not as clear since the action through the S3/S4 site tends to dominate and obscure events at this site.

Finally, binding to the top-VSD site presumably has complex effects (Fig. 7, yellow triangle), with interactions between the resin-acid derivative and R_2_ possibly stabilizing the C1 state, with R_3_ possibly stabilizing the O state and R_4_ possibly stabilizing O+ state. Thus its effect on the relative stabilization of the different states remains unclear.

### Quantitative evaluation of site-dependent contributions to the *G*(*V*) shifts

In an attempt to quantitatively evaluate the experimental data related to the suggested binding sites, we calculate here the contributions to the *G*(*V*) shifts due to binding either Wu50 or Wu161 to the various sites. We assume that (i) there is a background effect (probably due to binding to the VSD-top site, or to any other unidentified site), (ii) the compounds can bind independently to three sites: S3/S4, S4/pore, and “background” sites and that the contributions of the bound molecules to the *G*(*V*) shift are thus additive, and that (iii) there are four different binding contributions for the S4/pore site (each representing a mutation of this site). The assumption that binding to the sites is independent is in apparent contradiction to the idea that for Wu50, the 2R motif of the S3/S4 site overtrumps effects of the W454A mutation of the S4/pore site. However, as detailed below, for most mutants, a very good correlation between experimental data and model data is obtained without invoking interactions between the sites.

The expected shift for each mutant was

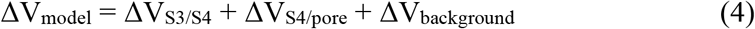

ΔV_S3/S4_ was set to 0 if the 2R motif was missing (i.e. in the WT channel). ΔV_S4/pore_ could adopt one of four values depending on the type of mutation (R362/W454 = WT configuration, R362Q, W454A, or R362Q/W454A). ΔV_background_ was assumed to be the same for all mutants. For Wu50 we have data for all eight possible combinations (8 mut; Supplementary Table S1). The best fit (least-square deviation between experimental and model data) was thus calculated for a system of eight equations. For Wu161, we have data for five of these mutations (5 mut). The lack of data for three of the mutations means fewer constraints during least squares fitting. To compare the fit for the two compounds in a fair manner, we also calculated the best solution for Wu50 using data for the five mutants available for Wu161 (Supplementary Table S1).

The correlation between experimental data and model data was good, supporting the assumption of independent binding sites (Supplementary Fig. S9A). It should be noted that the 8-mutant calculation for Wu50 showed the largest deviations for W454A and 2R/W454A, suggesting a putative interaction between these binding sites, as suggested previously. A summary of the results (Supplementary Fig. S9B) suggests the following:

- The background/residual effect was -8 mV (5 mut) to -10 mV (8 mut) for Wu50 (5 mut constrained to be the same as for 8 mut), while it was 0 to close to -50 mV for Wu161. A possible site of action for Wu50 is the VSD-top site. The data suggest that Wu161 may not bind here (or has no effect).
- The 2R motif (the S3/S4 site) contributed with -17 mV (8 mut) or -10 mV (5 mut) for Wu50, and with -17 mV (5 mutants) for Wu161. This suggests that Wu161 may have a larger effect than Wu50 when binding to this site with the 2R motif present.
- The WT-configuration of the S4/pore site (with a compound suggested to bind between R362 and W454 in the O state) contributes with -15 mV (8 mut) to -1820 mV (5 mut). Wu161 contributes with -8 to -132 mV. This suggests that Wu50 has a larger effect than Wu161, but despite the less rigid binding in MD, the effect is not absent.
- In the R362Q configuration of the S4/pore site, Wu50 is suggested to bind between W454 and R365 in a hypothetical O+ state. The contribution is very large for Wu50, -34 mV (8 mut) to -379 mV (5 mut). For Wu161 the contribution is only 0 to -6 mV.
- In the W454A configuration of the S4/pore site, the compounds are suggested to bind between R362 and F416 in the C1 state. Accordingly, the contributions of Wu50 and Wu161 are small (+20 to -6 mV; no clear difference between 5 mut and 8 mut).
- In the R362Q/W454A configuration of the S4/pore site, we only have data from the 8-mut model of Wu50. The compound is suggested to bind between F416 and R365 in the O state, and the contribution is very large, -31 mV

### The S3/S4 site

Both experimental data and MD simulations have previously been used to describe an interaction site for resin acids in the cleft between S3 and S4 (Fig 7, red) (Ottosson et al., 2017). We have also described that the 2R motif (M356R/A359R) (Fig 7, R_-1_, R_0_) greatly enhances the efficacy of resin acids (Ottosson et al., 2014, 2015, 2017; Silverå Ejneby et al., 2018). Here, we have shown that the 2R motif increases the Wu50-induced *G*(*V*) shift by 4-32 mV in the negative direction along the voltage axis, the magnitude of the shift depending on the background mutations. Binding to the S3/S4 site appears to be sensitive to the presence of charges at the extracellular end of S3 and S4. Some channels, such as K_V_2.1, have a native arginine at R_0_ (Supplementary Fig. S8). For Na_V_ and Ca_V_ channels (which are modulated by the resin acid, isopimaric acid; Salari et al., 2018), the charge profile varies from subunit to subunit. By designing resin-acid derivatives that primarily bind to the S3/S4 site, as appears to be the case for Wu161 and Wu181, it might be possible to engineer selectivity for specific channels or specific domains in heteromeric channels. As an example, resin-acid derivatives with a longer and flexible stalk are effective openers of the Shaker K_V_ channel with the 2R motif (Silverå Ejneby et al., 2018; present paper) and the K_V_7.2/7.3 channel (Silverå Ejneby et al., 2018). Thus, modifications of the stalk can be envisioned to obtain compounds selective for different K_V_ channel types.

### The S4/pore site

For the WT Shaker K_V_ channel, the S4/pore site, located between the VSD and the pore (Fig 7, purple), was found to be occupied by resin-acid derivatives. Binding to this site could possibly be disrupted by mutating W454 to alanine and/or by removing both R362 and R365 (Fig. 3) and was not as favorable for resin acids with a longer, flexible stalk. The presence of aromatic residues (W454 and/or F416 in Shaker) appeared important. A tryptophan in position 454 is unique for the Shaker K_V_ channel among all K_V_ channels (Supplementary Fig. S8), but K_V_3 and K_V_7 channels have a tryptophan in an adjacent position. F416, on the other hand, is either a phenylalanine or a tyrosine in most K_V_ channels and could potentially contribute to the general binding of resin-acid derivatives. Considering the presence of aromatic residues in various channels may be a way to engineer specificity for various channels (Supplementary Fig. S8).

### A promiscuous top-VSD site

Removing the 2R motif and disrupting the S4/pore site was however not enough to render the channel completely insensitive to Wu50. This residual *G(V)* shift could possibly depend on Wu50 binding to another site. Here, we have suggested that Wu50 can bind in a fairly promiscuous drug pocket located in a cleft in the top of the VSD in different voltage-gated ion channels (Fig. 7, yellow; Ahuja et al., 2015; Li et al., 2013; Liin et al., 2018a; Peretz et al., 2010; Li et al. 2020). From here the negatively charged Wu50 could contribute to the negative *G(V)* shift by electrostatically attracting S4 charges to rotate S4 in the clockwise direction and favor activation.

### Advantages and challenges of multi-site drug action

Traditional structure-based drug design aims to optimize compounds that fit in and strongly bind to a well-defined pocket of a biomolecule. This work, together with recent developments (Heusser et al., 2018), show that membrane proteins may be druggable via binding of compounds in a state-dependent manner, in different binding sites with similar binding affinity, and via mechanisms involving the membrane (Ahuja et al., 2015; Lambert et al., 2003; Wang, 2011). The resin-acid derivatives studied in this work seem to possess all of these three properties. These compounds seem to differentially stabilize the activated/open and the intermediate/closed states of the Shaker K_V_ channel depending on their binding to one or more of these three sites. Two of these sites are membrane facing, and binding of the compound to these sites involves displacing a previously bound lipid. While the lipid bilayer has so far often been considered an inert scaffold for membrane proteins, it is now recognized to often play an integral part in modulating protein function and in regulating the access of drug to proteins through competition with the compounds. This indicates that the membrane composition, known to vary from one cell type to the next, also offers possibilities for designing selective drugs with reduced off-target effects. For this to become a rational process, much of the molecular details underlying drug binding remain to be understood. We propose that studying the effects of resin-acid derivatives on a model channel constitute a first step in this direction. The experimental resolution of other channel states (intermediate and closed) will strengthen this type of work by enabling the explicit modeling and comparison of interactions of drugs with these other states.

Nevertheless, many fascinating questions remain after this work: for example, we were not able to assess binding affinity of the compounds to the different sites, such that we still do not know if the drugs bind in one or more sites at a time, or even if they bind to the same site in all four subunits. Finally, future research directions also involve deciphering the binding mechanism of the drugs to their site, particularly when it involves displacing lipids.

## ACKNOWLEDGEMENT

We thank Xiongyu Wu for synthesis of the compounds Wu50, Wu161, and Wu181 used in this study. We thank Antonios Pantazis for comments on the manuscript and Lea Rems for help with the revisions. This work was supported by grants from the Swedish Research Council, the Gustafsson Foundation and Science for Life Laboratory to LD, and the Swedish Research Council, the Swedish Brain Foundation, and the Swedish Heart-Lung Foundation to FE. The simulations were performed on resources provided by the Swedish National Infrastructure for Computing (SNIC) at PDC Centre for High Performance Computing (PDC-HPC).

## REFERENCES

Abraham, M.J., T. Murtola, R. Schulz, S. Páll, J.C. Smith, B. Hess, and E. Lindahl. 2015. GROMACS: High performance molecular simulations through multi-level parallelism from laptops to supercomputers. SoftwareX. 1–2:19–25. doi:10.1016/j.softx.2015.06.001.

Ahuja, S., S. Mukund, L. Deng, K. Khakh, E. Chang, H. Ho, S. Shriver, C. Young, S. Lin, J.P. Johnson, P. Wu, J. Li, M. Coons, C. Tam, B. Brillantes, H. Sampang, K. Mortara, K.K. Bowman, K.R. Clark, A. Estevez, Z. Xie, H. Verschoof, M. Grimwood, C. Dehnhardt, J.-C. Andrez, T. Focken, D.P. Sutherlin, B.S. Safina, M.A. Starovasnik, D.F. Ortwine, Y. Franke, C.J. Cohen, D.H. Hackos, C.M. Koth, and J. Payandeh. 2015. Structural basis of Nav1.7 inhibition by an isoform-selective small-molecule antagonist. Science. 350:aac5464. doi:10.1126/science.aac5464.

Beglov, D. and B. Roux. 1994. Finite representation of an infinite bulk system: Solvent boundary potential for computer simulations. J. Chem. Phys. 100:9050. doi: 10.1063/1.466711

Best, R. B., X. Zhu, J. Shim, P.E. Lopes, J. Mittal, M. Feig, A.D. Jr MacKerell. 2012. Optimization of the Additive CHARMM All-Atom Protein Force Field Targeting Improved Sampling of the Backbone ϕ, Ψ and Side-Chain χ1 and χ2 Dihedral Angles. J. Chem. Theory Comput. 8:3257–3273

Börjesson, S.I., and F. Elinder. 2011. An electrostatic potassium channel opener targeting the final voltage sensor transition. J Gen Physiol. 137:563–577. doi:10.1085/jgp.201110599.

Börjesson, S.I., T. Parkkari, S. Hammarström, and F. Elinder. 2010. Electrostatic Tuning of Cellular Excitability. Biophys J. 98:396–403. doi:10.1016/j.bpj.2009.10.026.

Brown, D.A., and P.R. Adams. 1980. Muscarinic suppression of a novel voltage-sensitive K+ current in a vertebrate neurone. Nature. 283:673–676.

Conti, L., J. Renhorn, A. Gabrielsson, F. Turesson, S.I. Liin, E. Lindahl, and F. Elinder. 2016. Reciprocal voltage sensor-to-pore coupling leads to potassium channel C-type inactivation. Sci Rep. 6:27562. doi:10.1038/srep27562.

Delemotte, L., M. Tarek, M.L. Klein, C. Amaral. W. Treptow. 2011 Intermediate states of the Kv1.2 voltage sensor from atomistic molecular dynamics simulations. Proc. Natl. Acad. Sci. USA 108:6109–6114

Henrion, U., J. Renhorn, S.I. Börjesson, E.M. Nelson, C.S. Schwaiger, P. Bjelkmar, B. Wallner, E. Lindahl, F. Elinder, 2012 Tracking a complete voltage-sensor cycle with metal-ion bridges. Proc. Natl. Acad. Sci. USA 109: 8552–8557

Heusser, S.A., M. Lycksell, X. Wang, S.E. McComas, R.J. Howard, and E. Lindahl. 2018. Allosteric potentiation of a ligand-gated ion channel is mediated by access to a deep membrane-facing cavity. PNAS. 115:10672–10677. doi:10.1073/pnas.1809650115.

Hoshi, T., W.N. Zagotta, and R.W. Aldrich. 1990. Biophysical and molecular mechanisms of Shaker potassium channel inactivation. Science. 250:533–538.

Humphrey, W., A. Dalke, and K. Schulten. 1996. VMD: Visual molecular dynamics. Journal of Molecular Graphics. 14:33–38. doi:10.1016/0263-7855(96)00018-5.

Imaizumi, Y., K. Sakamoto, A. Yamada, A. Hotta, S. Ohya, K. Muraki, M. Uchiyama, and T. Ohwada. 2002. Molecular Basis of Pimarane Compounds as Novel Activators of Large-Conductance Ca2+-Activated K+ Channel α-Subunit. Mol Pharmacol. 62:836–846. doi:10.1124/mol.62.4.836.

Jorgensen, W.L., J. Chandrasekhar, J.D. Madura, R.W. Impey, M.L. Klein. 1983. Comparison of simple potential functions for simulating liquid water. J. Chem. Phys. 79:926–935. Bibcode:1983JChPh..79..926J. doi:10.1063/1.445869.

Kamb, A., L.E. Iverson, and M.A. Tanouye. 1987. Molecular characterization of Shaker, a Drosophila gene that encodes a potassium channel. Cell. 50:405–413.

Kasimova, M.A., M. Tarek, A.K. Shaytan, K.V. Shaitan, and L. Delemotte. 2014. Voltage-gated ion channel modulation by lipids: Insights from molecular dynamics simulations. Biochimica et Biophysica Acta (BBA) - Biomembranes. 1838:1322–1331. doi:10.1016/j.bbamem.2014.01.024.

Klauda, J. B., R.M. Venable, J.A. Freites, J.W. O’Connor, D.J. Tobias, C. Mondragon-Ramirez, I. Vorobyov, A.D. Jr MacKerel, R.W. Pastor. 2010. Update of the CHARMM all-atom additive force field for lipids: validation on six lipid types. J. Phys. Chem. B 114:7830–7843

Kobayashi, K., Y. Nishizawa, K. Sawada, H. Ogura, and M. Miyabe. 2008. K(+)-channel openers suppress epileptiform activities induced by 4-aminopyridine in cultured rat hippocampal neurons. J. Pharmacol. Sci. 108:517–528.

Lambert, J.J., D. Belelli, D.R. Peden, A.W. Vardy, and J.A. Peters. 2003. Neurosteroid modulation of GABAA receptors. Prog. Neurobiol. 71:67–80. doi:10.1016/j.pneurobio.2003.09.001.

Lee, J., X. Cheng, J.M. Swails, M.S. Yeom, P.K. Eastman, J.A. Lemkul, S. Wei, J. Buckner, J.C. Jeong, Y. Qi, S. Jo, V.S. Pande, D.A. Case, C.L. Brooks, A.D. MacKerell, J.B. Klauda, and W. Im. 2016. CHARMM-GUI Input Generator for NAMD, GROMACS, AMBER, OpenMM, and CHARMM/OpenMM Simulations Using the CHARMM36 Additive Force Field. J. Chem. Theory Comput. 12:405–413. doi:10.1021/acs.jctc.5b00935.

Li, P., Z. Chen, H. Xu, H. Sun, H. Li, H. Liu, H. Yang, Z. Gao, H. Jiang, and M. Li. 2013. The gating charge pathway of an epilepsy-associated potassium channel accommodates chemical ligands. Cell Res. 23:1106–1118. doi:10.1038/cr.2013.82.

Li X, Zhang Q, Guo P, Fu J, Mei L, Lv D, et al. 2020. Molecular basis for ligand activation of the human KCNQ2 channel. Cell Res. doi:10.1038/s41422-020-00410-8

Liin, S.I., P.-E. Lund, J.E. Larsson, J. Brask, B. Wallner, and F. Elinder. 2018a. Biaryl sulfonamide motifs up- or down-regulate ion channel activity by activating voltage sensors. J. Gen. Physiol. 150:1215–1230. doi:10.1085/jgp.201711942.

Liin, S.I., S. Yazdi, R. Ramentol, R. Barro-Soria, and H.P. Larsson. 2018b. Mechanisms Underlying the Dual Effect of Polyunsaturated Fatty Acid Analogs on Kv7.1. Cell Rep. 24:2908–2918. doi:10.1016/j.celrep.2018.08.031.

McGibbon, R.T., K.A. Beauchamp, M.P. Harrigan, C. Klein, J.M. Swails, C.X. Hernández, C.R. Schwantes, L.-P. Wang, T.J. Lane, and V.S. Pande. 2015. MDTraj: A Modern Open Library for the Analysis of Molecular Dynamics Trajectories. Biophys J. 109:1528–1532. doi:10.1016/j.bpj.2015.08.015.

Ottosson, N.E., S.I. Liin, and F. Elinder. 2014. Drug-induced ion channel opening tuned by the voltage sensor charge profile. J Gen Physiol. 143:173–182. doi:10.1085/jgp.201311087.

Ottosson, N.E., M. Silverå Ejneby, X. Wu, S. Yazdi, P. Konradsson, E. Lindahl, and F. Elinder. 2017. A drug pocket at the lipid bilayer-potassium channel interface. Sci Adv. 3:e1701099. doi:10.1126/sciadv.1701099.

Ottosson, N.E., X. Wu, A. Nolting, U. Karlsson, P.-E. Lund, K. Ruda, S. Svensson, P. Konradsson, and F. Elinder. 2015. Resin-acid derivatives as potent electrostatic openers of voltage-gated K channels and suppressors of neuronal excitability. Sci Rep. 5:13278. doi:10.1038/srep13278.

Papazian, D.M., X.M. Shao, S.A. Seoh, A.F. Mock, Y. Huang, and D.H. Wainstock. 1995. Electrostatic interactions of S4 voltage sensor in Shaker K+ channel. Neuron. 14:1293–1301.

Peretz, A., L. Pell, Y. Gofman, Y. Haitin, L. Shamgar, E. Patrich, P. Kornilov, O. Gourgy-Hacohen, N. Ben-Tal, and B. Attali. 2010. Targeting the voltage sensor of Kv7.2 voltage-gated K+ channels with a new gating-modifier. Proc. Natl. Acad. Sci. U.S.A. 107:15637–15642. doi:10.1073/pnas.0911294107.

Sakamoto, K., T. Nonomura, S. Ohya, K. Muraki, T. Ohwada, and Y. Imaizumi. 2006. Molecular Mechanisms for Large Conductance Ca2+-Activated K+ Channel Activation by a Novel Opener, 12,14-Dichlorodehydroabietic Acid. J Pharmacol Exp Ther. 316:144–153. doi:10.1124/jpet.105.093856.

Sakamoto, K., Y. Suzuki, H. Yamamura, S. Ohya, K. Muraki, and Y. Imaizumi. 2017. Molecular mechanisms underlying pimaric acid-induced modulation of voltage-gated K+ channels. Journal of Pharmacological Sciences. 133:223–231. doi:10.1016/j.jphs.2017.02.013.

Salari, S., M. Silverå Ejneby, J. Brask, and F. Elinder. 2018. Isopimaric acid - a multi-targeting ion channel modulator reducing excitability and arrhythmicity in a spontaneously beating mouse atrial cell line. Acta Physiol (Oxf). 222. doi:10.1111/apha.12895.

Silverå Ejneby, M., X. Wu, N.E. Ottosson, E.P. Münger, I. Lundström, P. Konradsson, and F. Elinder. 2018. Atom-by-atom tuning of the electrostatic potassium-channel modulator dehydroabietic acid. J. Gen. Physiol. 150:731–750. doi:10.1085/jgp.201711965.

Smith-Maxwell, C.J., J.L. Ledwell, and R.W. Aldrich. 1998. Uncharged S4 residues and cooperativity in voltage-dependent potassium channel activation. J. Gen. Physiol. 111:421–439.

Tombola, F., M.M. Pathak, P. Gorostiza, and E.Y. Isacoff. 2007. The twisted ion-permeation pathway of a resting voltage-sensing domain. Nature. 445:546–549. doi: 10.1038/nature05396.

Trott, O., and A.J. Olson. 2010. AutoDock Vina: improving the speed and accuracy of docking with a new scoring function, efficient optimization and multithreading. J Comput Chem. 31:455–461. doi:10.1002/jcc.21334.

Vanommeslaeghe, K., E. Hatcher, C. Acharya, S. Kundu, S. Zhong, J. Shim, E. Darian, O. Guvench, P. Lopes, I. Vorobyov A.D. Jr MacKerell 2010. CHARMM general force field: A force field for drug-like molecules compatible with the CHARMM all-atom additive biological force fields. J. Comput. Chem. 31:671–690

Wang, H.-S., Z. Pan, W. Shi, B.S. Brown, R.S. Wymore, I.S. Cohen, J.E. Dixon, and D. McKinnon. 1998. KCNQ2 and KCNQ3 Potassium Channel Subunits: Molecular Correlates of the M-Channel. Science. 282:1890–1893. doi:10.1126/science.282.5395.1890.

Wang, M. 2011. Neurosteroids and GABA-A Receptor Function. Front Endocrinol (Lausanne). 2:44. doi:10.3389/fendo.2011.00044.

Wu, C., K. V Gopal, T.J. Lukas, G.W. Gross, and E.J. Moore. 2014a. Pharmacodynamics of potassium channel openers in cultured neuronal networks. Eur. J. Pharmacol. 732:68–75. doi:10.1016/j.ejphar.2014.03.017.

Wu, E.L., X. Cheng, S. Jo, H. Rui, K.C. Song, E.M. Dávila-Contreras, Y. Qi, J. Lee, V. Monje-Galvan, R.M. Venable, J.B. Klauda, and W. Im. 2014b. CHARMM-GUI Membrane Builder Toward Realistic Biological Membrane Simulations. J Comput Chem. 35:1997–2004. doi:10.1002/jcc.23702.

Yazdi, S., M. Stein, F. Elinder, M. Andersson, and E. Lindahl. 2016. The Molecular Basis of Polyunsaturated Fatty Acid Interactions with the Shaker Voltage-Gated Potassium Channel. PLoS Comput. Biol. 12:e1004704. doi:10.1371/journal.pcbi.1004704.

Zoete, V., M.A. Cuendet, A. Grosdidier, and O. Michielin. 2011. SwissParam: A fast force field generation tool for small organic molecules. Journal of Computational Chemistry. 32:2359–2368. doi:10.1002/jcc.21816.

