## Supplementary Figures and Text for "Resin-acid derivatives bind to multiple sites on the voltage-sensor domain of the Saker channel"

Supplementary Figures S1-S9
Supplementary Table S1
Supplementary Text (including Figures S10-S11): The role of coupling between early and late voltage-sensor transitions for the interpretation of *G*(*V*) shifts

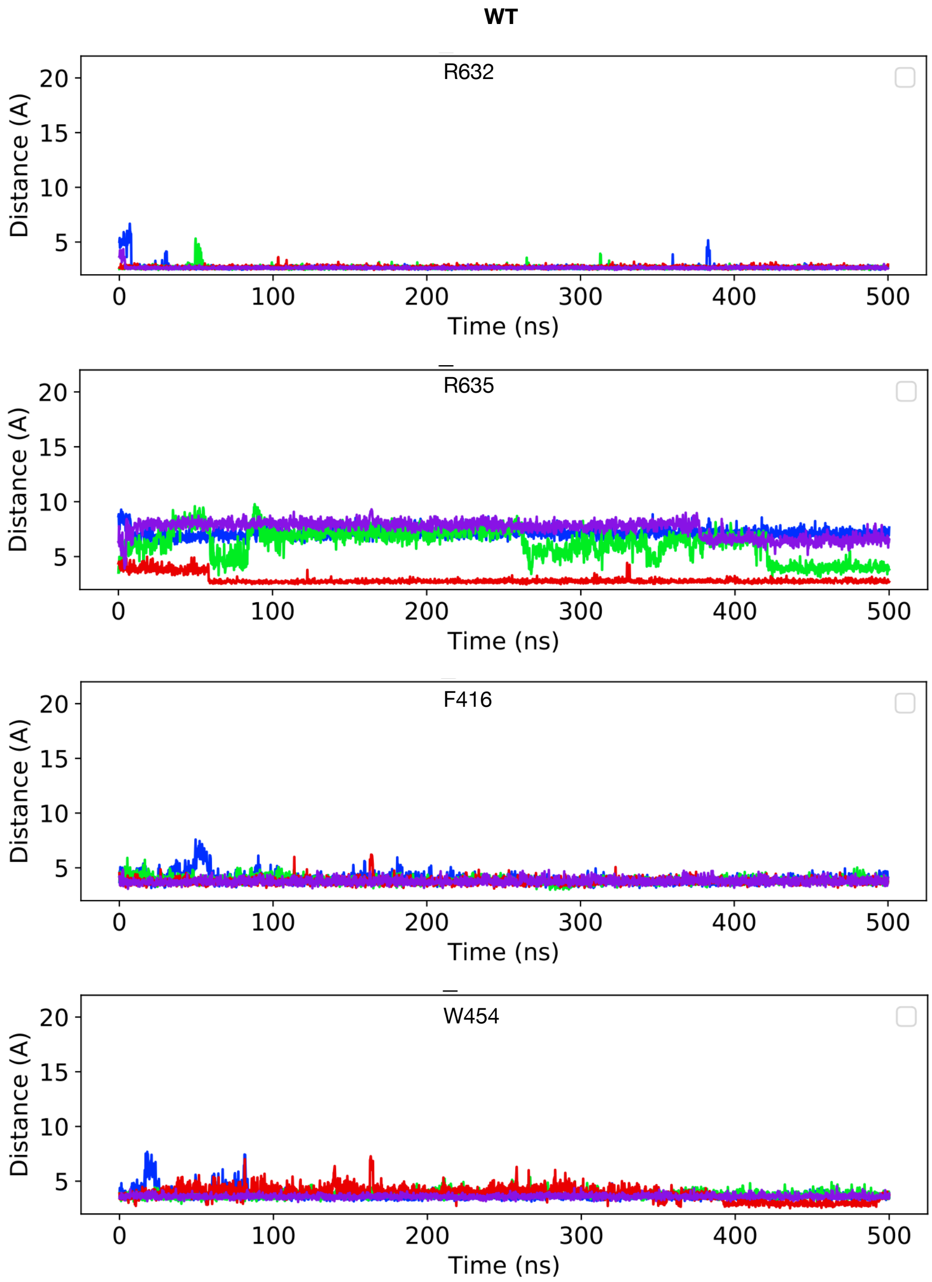

**Figure S1** Distance between closest atoms of Wu50 and R362, R365, F416 and W454 along the MD simulations of the WT channel system. Each of four subunits is depicted in a different color.

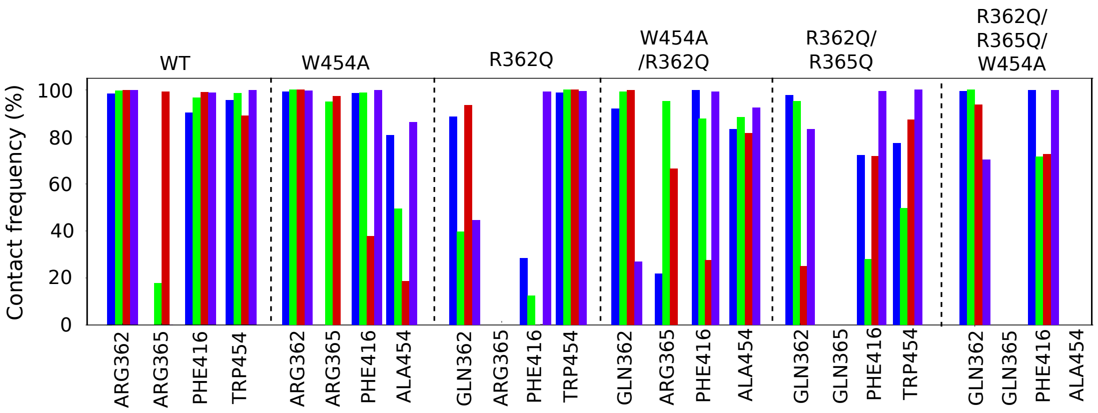

**Figure S2** Contact frequency between any atom of Wu50 and select S4/pore site residues in the WT, W454A, R362Q, W454A/R362Q, R362Q/R365Q, W454A/R362Q/R365Q channel simulations. Each of four subunits is depicted in a different color, following the color scheme presented in Figure 1.

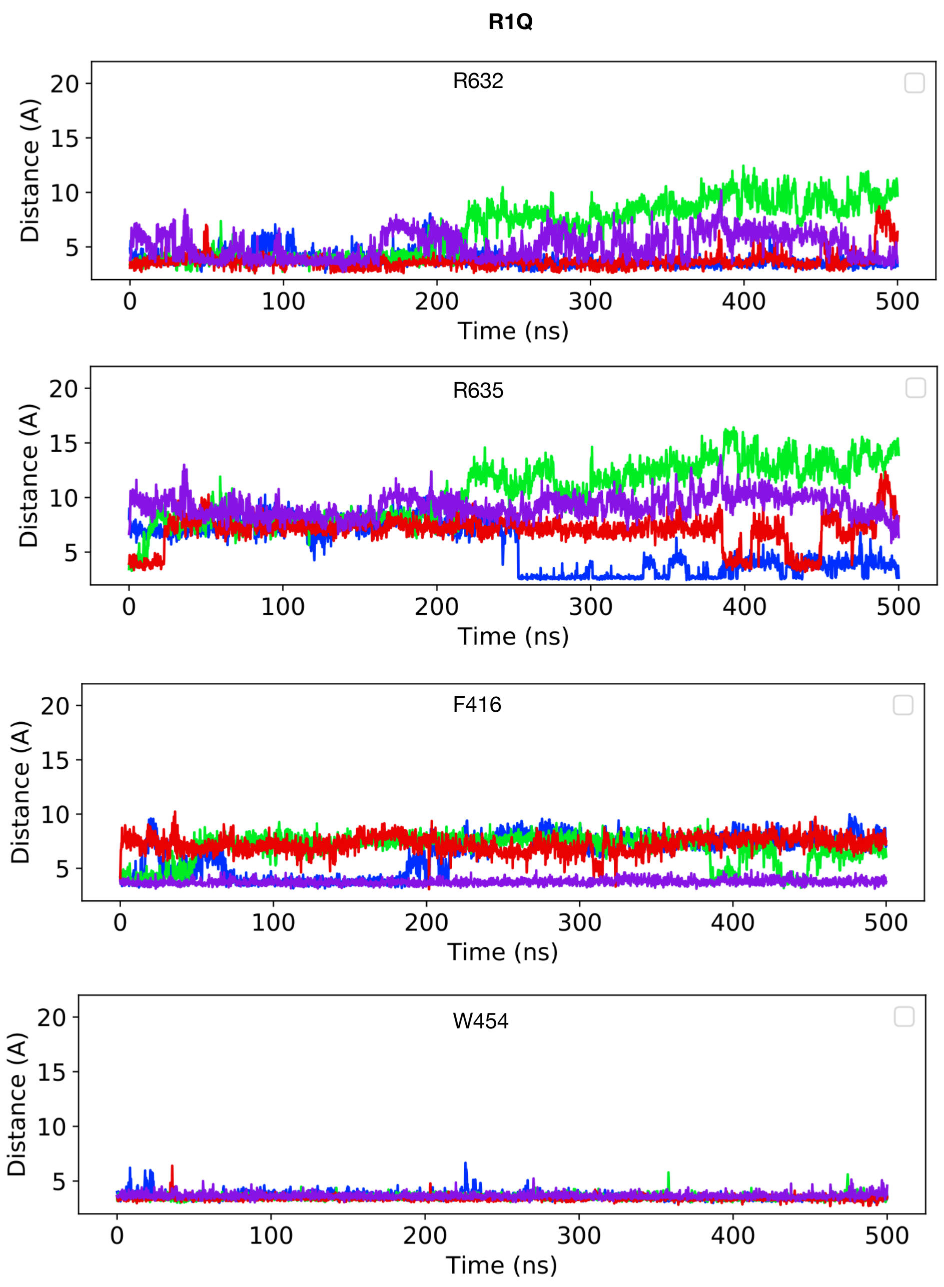

**Figure S3** Distance between closest atoms of Wu50 and R362, R365, F416 and W454 along the MD simulations of the R362Q channel system. Each of four subunits is depicted in a different color.

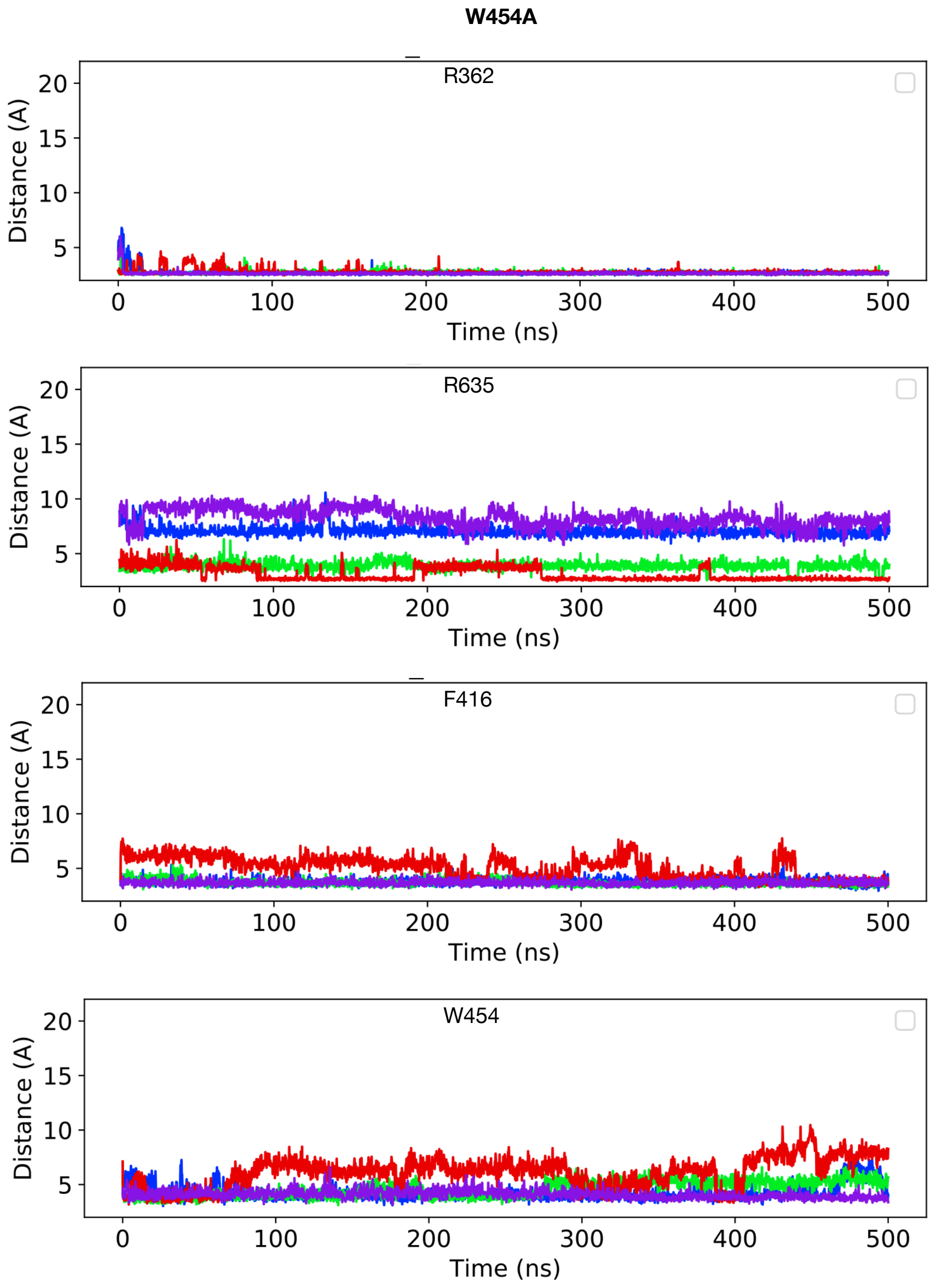

**Figure S4** Distance between closest atoms of Wu50 and R362, R365, F416 and W454 along the MD simulations of the W454A channel system. Each of four subunits is depicted in a different color.

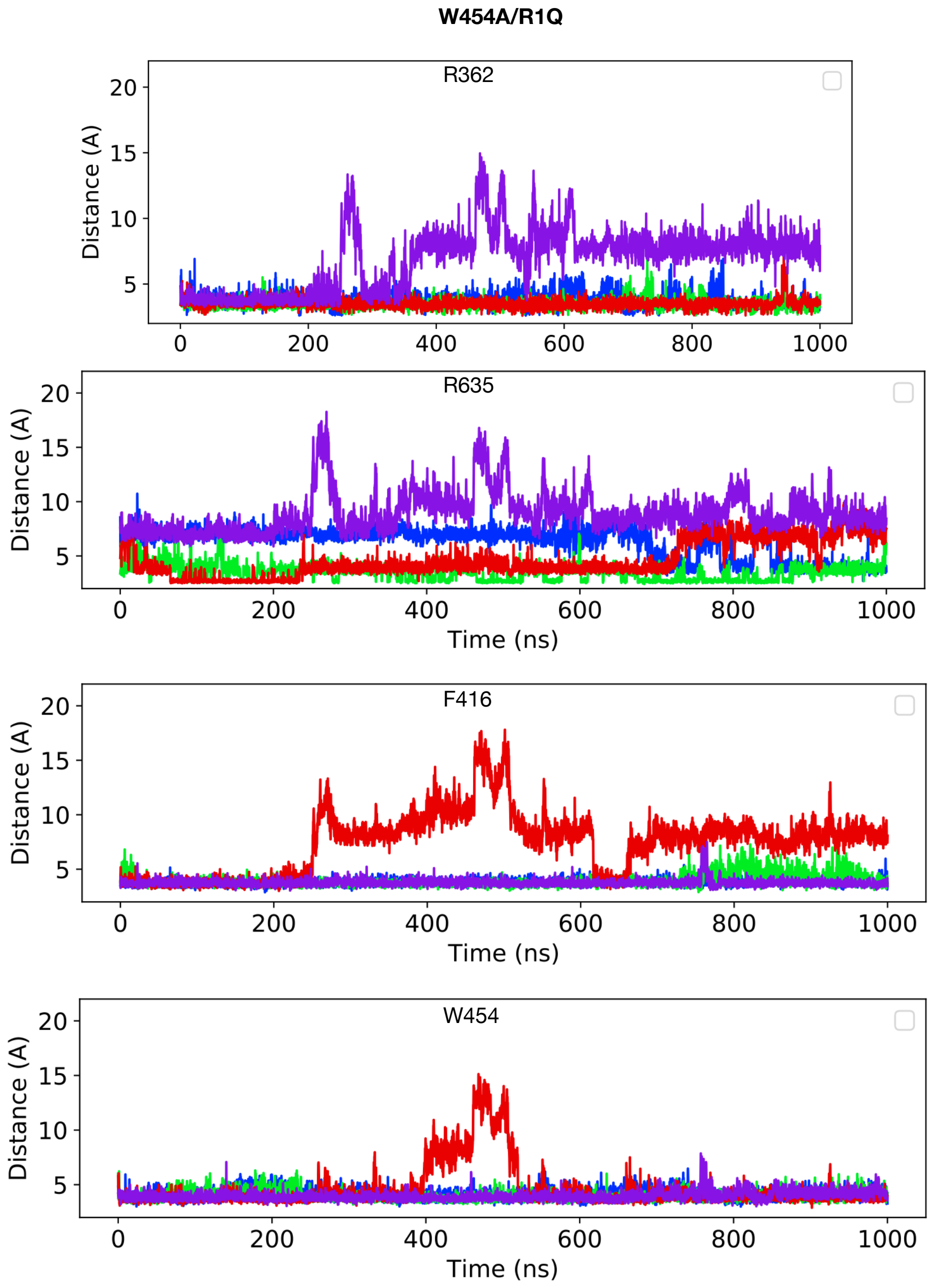

**Figure S5** Distance between closest atoms of Wu50 and R362, R365, F416 and W454 along the MD simulations of the W454A/R362Q channel system. Each of four subunits is depicted in a different color.

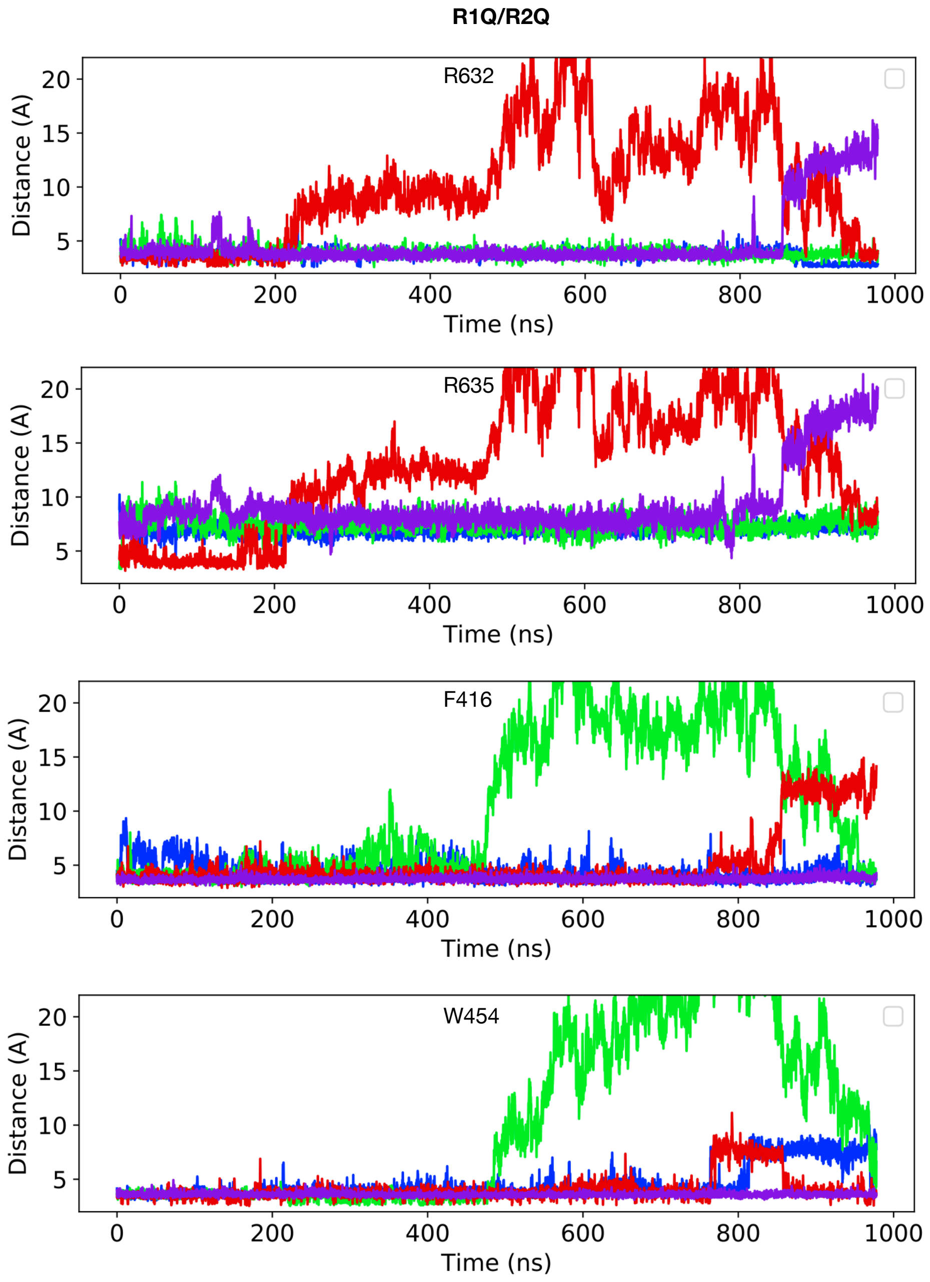

**Figure S6** Distance between closest atoms of Wu50 and R362, R365, F416 and W454 along the MD simulations of the R362Q/R365Q channel system. Each of four subunits is depicted in a different color.

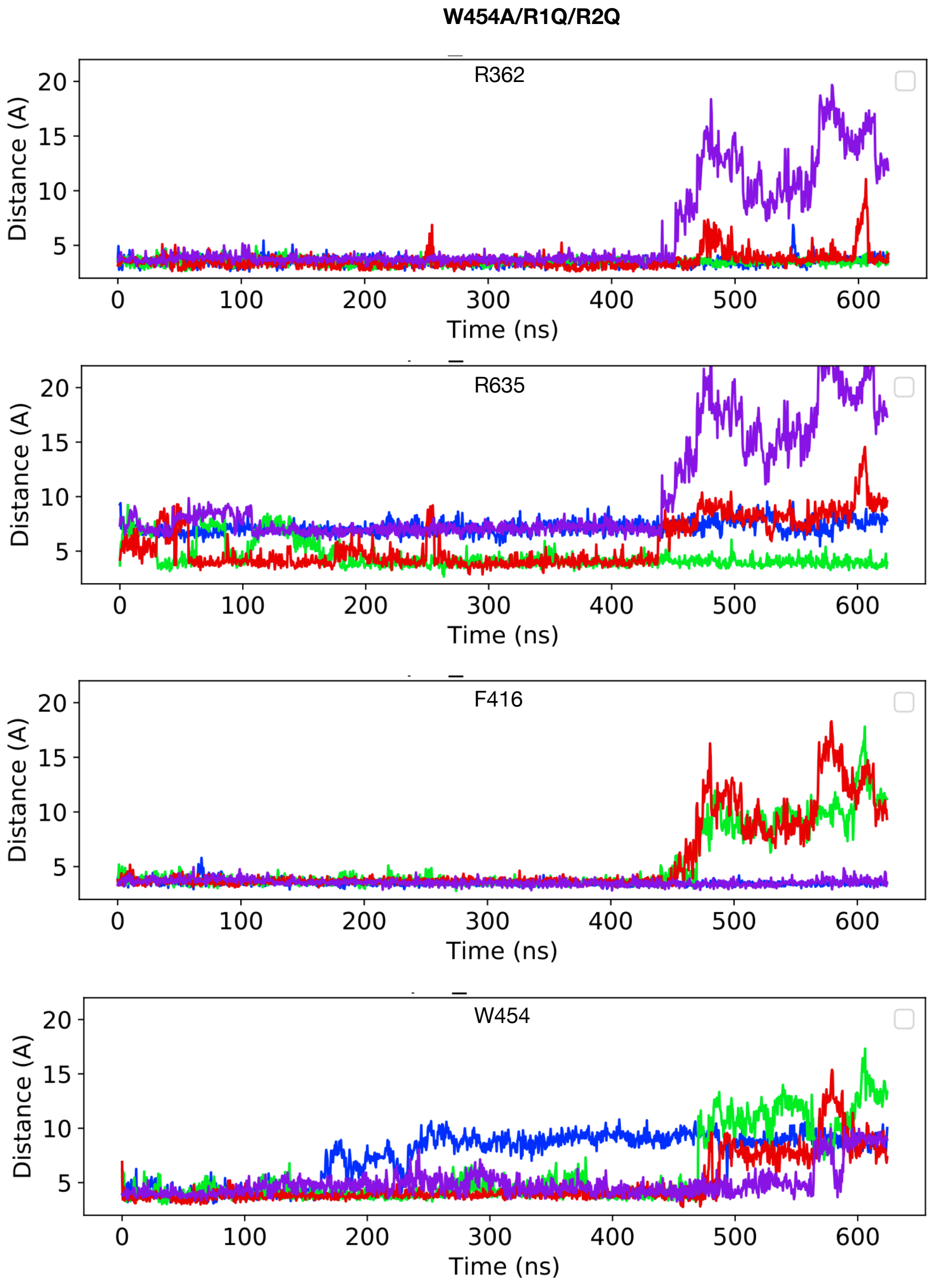

**Figure S7** Distance between closest atoms of Wu50 and R362, R365, F416 and W454 along the MD simulations of the R362Q/R365Q/W454A channel system. Each of four subunits is depicted in a different color.

```
              e.c.     S4     i.c.    i.c.       S5       e.c.       e.c.              S6              i.c.

        356359362365                                       416         454
         ↓   ↓   ↓   ↓                                      ↓           ↓
Shaker   MSLAILRVIRLVRVFRIFKLSRH     RELGLLIFFLFIGVVLFSSAVYFAEA    PVGVWGKIVGSLCAIAGVLTIALPVPVIVSNFNYFYHRET
hKv1.1   TSLAILRVIRLVRVFRIFKLSRH     RELGLLIFFLFIGVILFSSAVYFAEA    PVTIGGKIVGSLCAIAGVLTIALPVPVIVSNFNYFYHRET
hKv1.2   MSLAILRVIRLVRVFRIFKLSRH     RELGLLIFFLFIGVILFSSAVYFAEA    PTTIGGKIVGSLCAIAGVLTIALPVPVIVSNFNYFYHRET
hKv1.3   MSLAILRVIRLVRVFRIFKLSRH     RELGLLIFFLFIGVILFSSAVYFAEA    PVTIGGKIVGSLCAIAGVLTIALPVPVIVSNFNYFYHRET
hKv1.4   MSFAILRIIRLVRVFRIFKLSRH     RELGLLIFFLFIGVILFSSAVYFAEA    PITVGGKIVGSLCAIAGVLTIALPVPVIVSNFNYFYHRET
hKv1.5   MSLAILRVIRLVRVFRIFKLSRH     RELGLLIFFLFIGVILFSSAVYFAEA    PITVGGKIVGSLCAIAGVLTIALPVPVIVSNFNYFYHRET
hKv1.6   MSLAILRVIRLVRVFRIFKLSRH     RELGLLIFFLFIGVILFSSAVYFAEA    PMTVGGKIVGSLCAIAGVLTIALPVPVIVSNFNYFYHRET
hKv1.7   MSLAILRVIRLVRVFRIFKLSRH     RELGLLIFFLFIGVVLFSSAVYFAEV    PVTVGGKIVGSLCAIAGVLTISLPVPVIVSNFSYFYHRET
hKv1.8   MSLAILRIIRLVRVFRIFKLSRH     RELGLLIFFLFIGVILFSSAVYFAEV    PTTPGGKIVGTLCAIAGVLTIALPVPVIVSNFNYFYHRET

hKv2.1   NVRRVVQIFRIMRILRILKLARH     NELGLLILFLAMGIMIFSSLVFFAEK    PKTLLGKIVGGLCCIAGVLVIALPIPIIVNNFSEFYKEQK
hKv2.2   NVRRVVQIFRIMRILRILKLARH     NELGLLILFLAMGIMIFSSLVFFAEK    PKTLLGKIVGGLCCIAGVLVIALPIPIIVNNFSEFYKEQK

hKv3.1   DVLGFLRVVRFVRILRIFKLTRH     NEFLLLIIFLALGVLIFATMIYYAER    PQTWSGMLVGALCALAGVLTIAMPVPVIVNNFGMYYSLAM
hKv3.2   DVLGFLRVVRFVRILRIFKLTRH     NEFLLLIIFLALGVLIFATMIYYAER    PQTWSGMLVGALCALAGVLTIAMPVPVIVNNFGMYYSLAM
hKv3.3   DVLGFLRVVRFVRILRIFKLTRH     NEFLLLIIFLALGVLIFATMIYYAER    PKTWSGMLVGALCALAGVLTIAMPVPVIVNNFGMYYSLAM
hKv3.4   DVLGFLRVVRFVRILRIFKLTRH     NEFLLLIIFLALGVLIFATMIYYAER    PKTWSGMLVGALCALAGVLTIAMPVPVIVNNFGMYYSLAM

hKv4.1   DVSGAFVTLRVFRVFRIFKFSRH     SELGFLLFSLTMAIIIFATVMFYAEK    PSTIAGKIFGSICSLSGVLVIALPVPVIVSNFSRIYHQNQ
hKv4.2   DVSGAFVTLRVFRVFRIFKFSRH     SELGFLLFSLTMAIIIFATVMFYAEK    PKTIAGKIFGSICSLSGVLVIALPVPVIVSNFSRIYHQNQ
hKv4.3   DVSGAFVTLRVFRVFRIFKFSRH     SELGFLLFSLTMAIIIFATVMFYAEK    PKTIAGKIFGSICSLSGVLVIALPVPVIVSNFSRIYHQNQ

hKv7.1   FATSAIRGIRFLQILRMLHVDRQ     QELITTLYIGFLGLIFSSYFVYLAEK    PQTWVGKTIASCFSVFAISFFALPAGILGSGFALKVQQKQ
hKv7.2   FATSALRSLRFLQILRMIRMDRR     KELVTAWYIGFLCLILASFLVYLAEK    PQTWNGRLLAATFTLIGVSFFALPAGILGSGFALKVQEQH
hKv7.3   LATS LRSLRFLQILRMLRMDRR     KELITAWYIGFLTLILSSFLVYLVEK    PKTWEGRLIAATFSLIGVSFFALPAGILGSGLALKVQEQH
hKv7.4   FATSALRSMRFLQILRMVRMDRR     KELITAWYIGFLVLIFASFLVYLAEK    PHTWLGRVLAAGFALLGISFFALPAGILGSGFALKVQEQH
hKv7.5   FATSALRSLRFLQILRMVRMDRR     KELITAWYIGFLVLIFSSFLVYLVEK    PLTWLGRLLSAGFALLGISFFALPAGILGSGFALKVQEQH
```

**Figure S8** Sequence alignment between segments S4, S5 and S6 in several $K_V$ channel families, highlighting positively and negatively charged residues (blue and red, respectively) as well as aromatic ones (green). Gray bars are approximate transmembrane helical segments. E.c. = extracellular. i.c. = intracellular. Arrows denote mutated residues in the present investigation.

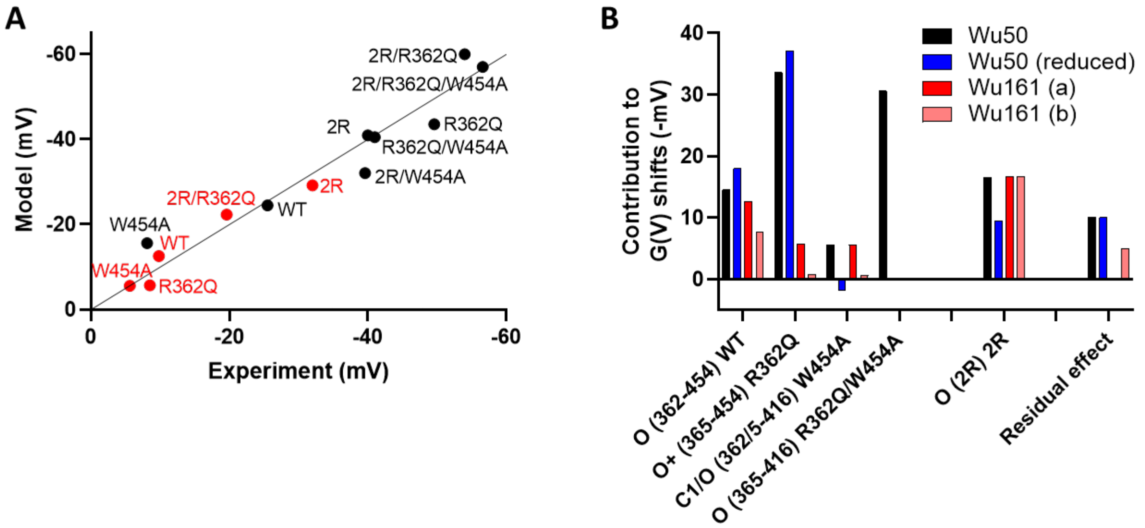

**Figure S9 A)** Correlation between experimental data and model data as described in the Discussion of the main text. Color coding in B. **B)** Best solutions to the models described in the Discussion of the main text. Wu50 data is based on eight mutants. Wu50 (reduced) data is based on a reduced set of five mutations (residual effect fixed to black bar). Wu161 data is based on five mutants (a-variant, residual effect = 0; b-variant, largest residual effect with no bars below zero-line). For O (365-416) R362Q/W454A only the black bar exists.

### SUPPLEMENTARY TABLES

### Table S1: Summary of *G*(*V*) shifts and $G_{MAX}$ effects induced by 100 µM of either Wu50 or Wu161 for the mutants reported in this paper

| | Wu50 pH 9.0 | | | | | | Wu161 pH 7.4 | | | | | |
|---|---|---|---|---|---|---|---|---|---|---|---|---|
| **Mutant** | **ΔVG(V) (mV)** | | | **$G_{MAX}$ (rel.)** | | | **ΔVG(V) (mV)** | | | **Gmax (rel.)** | | |
| | Mean | SEM | n | **Mean** | SEM | n | Mean | SEM | n | **Mean** | SEM | n |
| **M356R/A359R (=2R)** | -40.0 | 2.7 | 10 | 1.58 | 0.09 | 8 | -32.0 | 2.7 | 6 | 1.20 | 0.05 | 6 |
| **M356R/A359R/W454A** | -39.6 | 2.9 | 5 | 1.28 | 0.09 | 5 | | | | | | |
| **M356R/A359R/R362Q** | -54.0 | 2.3 | 4 | 1.93 | 0.25 | 4 | -19.6 | 4.1 | 5 | 1.19 | 0.07 | 5 |
| **M356R/A359R/R362Q/W454A** | -56.6 | 5.8 | 4 | 1.50 | 0.14 | 4 | | | | | | |
| **WT** | -25.5 | 1.4 | 5 | 1.36 | 0.07 | 6 | -9.8 | 1.1 | 5 | 1.03 | 0.04 | 3 |
| **W454A** | -8.1 | 0.4 | 5 | 0.56 | 0.08 | 5 | -5.6 | 0.8 | 4 | 0.86 | 0.04 | 4 |
| **R362Q** | -49.6 | 1.2 | 3 | 1.25 | 0.14 | 3 | -8.5 | 0.5 | 3 | 1.05 | 0.01 | 4 |
| **R362Q/W454A** | -41.0 | 4.4 | 5 | 1.28 | 0.08 | 5 | | | | | | |
| **R362Q/W454A/F416A** | -34.3 | 3.4 | 3 | 0.17 | 0.06 | 5 | 2.0 | 1.4 | 4 | 0.47 | 0.07 | 4 |
| **R362Q/R365Q** | -25.5 | 4.6 | 5 | 0.55 | 0.12 | 5 | -5.2 | 1.2 | 4 | 0.99 | 0.03 | 4 |
| **R362Q/R365Q/W454A** | -24.7 | 3.0 | 5 | 0.57 | 0.13 | 6 | | | | | | |
| **R362Q/R365Q/W454A/F280I** | ND | N/A | N/A | 0.05 | 0.02 | 4 | 1.1 | 3.1 | 3 | 0.64 | 0.12 | 3 |

**The role of coupling between early and late voltage-sensor transitions for the interpretation of *G*(*V)* shifts**

Opening of voltage-gated ion channels is controlled by the movement of voltage sensors. Several studies suggest that this channel opening is controlled by at least two types of voltage-dependent transitions, sometimes called Q1 and Q2 (Zagotta et al., 1994; Keynes and Bezanilla et al., 1994; Zagotta et al., 1994; Schoppa and Sigworth, 1998; Elinder, 1998; Baker et al., 1998). When the membrane potential is changed from a normal resting potential, where the channel is closed, to a more positive voltage where the channel is open, the gating charge(s) Q1 will move first, followed by the gating charge(s) Q2.

In general, Q1 and Q2 are tightly coupled to each other, meaning that they occur at approximately the same voltage and thus both are tightly coupled to the channel opening. However, some mutations separate Q1 and Q2 from each other, so that channel opening is only controlled by Q2 (Schoppa et al., 1992; Smith-Maxwell et al., 1998). This separation shifts channel opening (*G(V)*) to more positive voltages than the voltage where the majority of the gating charges (Q1) move (Q(V)); these mutations also makes the *G(V)* shallower than in wild-type.

In previous studies, we have suggested that polyunsaturated fatty acids and resin acids almost exclusively act on the last voltage sensor transition, associated with Q2, to shift *G(V)* (Börjesson and Elinder, 2011; Ottosson et al., 2017). Thus, the ILT-mutant (Smith-Maxwell et al., 1998), which separates Q1 and Q2 from each other by 150-200 mV, uncovers the effect of the compounds; while in wild-type the effect on the final, channel-opening step, is largely masked by the tight coupling between Q1 and Q2.

To model this effect we have developed a relatively simple but robust steady-state gating model (Börjesson & Elinder, 2011). In the wild-type model, the Q1 and Q2 transitions completely overlap (0 mV shift in the x-direction in Fig. S10); the midpoint of the *G(V)* curve is 20 mV more positive than the midpoints of Q1 and Q2. In the ILT mutant, which has a Q1-Q2 separation of 180 mV, the midpoint of the *G(V)* curve is located +180 mV relative to Q1 (Fig. S10).

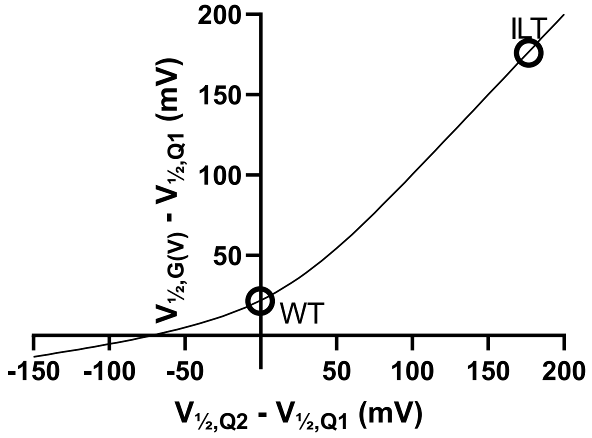

Fig. S10. Difference between the *G(V)* midpoint and the Q1 midpoint plotted against the difference between the Q2 and Q1 midpoints.

When Q1 and Q2 are separated the *G(V)* curve becomes shallower (Fig. S11), corresponding to a slope value increase (*G(V)* data is fitted to $G(V) = 1 / (1+\exp(-(V-V_{½})/s))$, where V is the membrane potential, $V_{½}$ the midpoint of the curve, and s is the slope value).

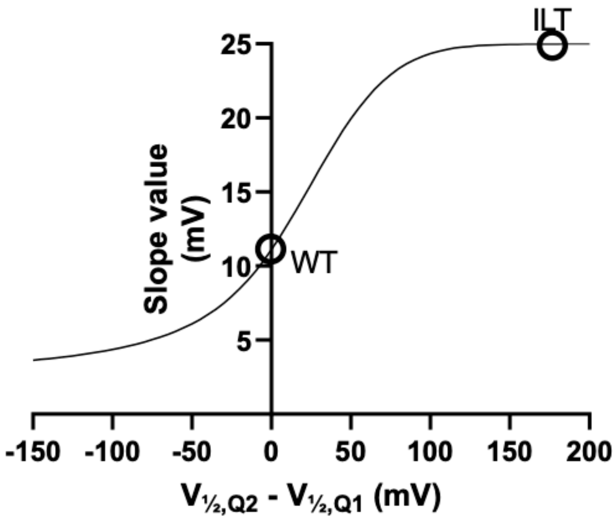

Fig. S11. *G(V)* slope plotted against the difference between the Q2 and Q1 midpoints.

Thus, if a mutation separates Q2 from Q1, we would expect a shallower *G(V)* curve shifted in positive direction along the voltage axis (if Q1 is not shifted). If a compound exclusively acts on the opening step (Q2) then we would expect an increase in *G(V)* shift compared with wild-type.

So, what about the R362Q mutation? The *G(V)* is shifted +25 mV compared with wild-type (Ottosson et al., 2014). The *Q(V)* is shifted approximately +10 to +15 mV (Seoh et al., 1996). Thus the Q1/Q2 separation is about 10-15 mV. This suggests that the mutations should make the *G(V)* slightly shallower (slope factor increase about 20%), which is consistent with our experimental data. This separation also suggests that the compound-induced *G(V)* shift should increase by approximately 20% (can be derived from the slope at 0 mV and +10 to +15 mV of the curve in Fig. S10). Thus, the R362Q mutation is expected to increase the *G(V)* shift from -25.5 mV to approximately -30 mV. Experimentally we found a shift of -49.6 mV suggesting that a very large part of the effect is caused by direct effects via the site of action and not by an indirect gating effect such as the one described here.

**References**

Baker OS, Larsson HP, Mannuzzu LM, Isacoff EY. Three transmembrane conformations and sequence-dependent displacement of the S4 domain in shaker K+ channel gating. Neuron. 1998 Jun;20(6):1283-94. doi: 10.1016/s0896-6273(00)80507-3.

Bezanilla F, Perozo E, Stefani E. Gating of Shaker K+ channels: II. The components of gating currents and a model of channel activation. Biophys J. 1994 Apr;66(4):1011-21. doi: 10.1016/S0006-3495(94)80882-3.

Börjesson SI, Elinder F. An electrostatic potassium channel opener targeting the final voltage sensor transition. J Gen Physiol. 2011 Jun;137(6):563-77. doi: 10.1085/jgp.201110599.

Keynes RD, Elinder F. Modelling the activation, opening, inactivation and reopening of the voltage-gated sodium channel. Proc Biol Sci. 1998 Feb 22;265(1393):263-70. doi: 10.1098/rspb.1998.0291.

Ottosson NE, Liin SI, Elinder F. Drug-induced ion channel opening tuned by the voltage sensor charge profile. J Gen Physiol. 2014 Feb;143(2):173-82. doi: 10.1085/jgp.201311087. Epub 2014 Jan 13.

Ottosson NE, Silverå Ejneby M, Wu X, Yazdi S, Konradsson P, Lindahl E, Elinder F. A drug pocket at the lipid bilayer-potassium channel interface. Sci Adv. 2017 Oct 25;3(10):e1701099. doi: 10.1126/sciadv.1701099. eCollection 2017 Oct.

Schoppa NE, McCormack K, Tanouye MA, Sigworth FJ. The size of gating charge in wild-type and mutant Shaker potassium channels. Science. 1992 Mar 27;255(5052):1712-5. doi: 10.1126/science.1553560.

Schoppa NE, Sigworth FJ. Activation of Shaker potassium channels. III. An activation gating model for wild-type and V2 mutant channels. J Gen Physiol. 1998 Feb;111(2):313-42. doi: 10.1085/jgp.111.2.313.

Seoh SA, Sigg D, Papazian DM, Bezanilla F. Voltage-sensing residues in the S2 and S4 segments of the Shaker K+ channel. Neuron. 1996 Jun;16(6):1159-67. doi: 10.1016/s0896-6273(00)80142-7.

Smith-Maxwell CJ, Ledwell JL, Aldrich RW. Uncharged S4 residues and cooperativity in voltage-dependent potassium channel activation. J Gen Physiol. 1998 Mar;111(3):421-39. doi: 10.1085/jgp.111.3.421.

Zagotta WN, Hoshi T, Aldrich RW. Shaker potassium channel gating. III: Evaluation of kinetic models for activation. J Gen Physiol. 1994 Feb;103(2):321-62. doi: 10.1085/jgp.103.2.321.
